# Spt4 Facilitates the Movement of RNA Polymerase II through the +2 Nucleosomal Barrier

**DOI:** 10.1101/2021.03.03.433772

**Authors:** Ülkü Uzun, Thomas Brown, Harry Fischl, Andrew Angel, Jane Mellor

**Affiliations:** Department of Biochemistry, University of Oxford, South Parks Road, Oxford OX1 3QU, UK

## Abstract

Spt4 is a transcription elongation factor, with homologues in organisms with nucleosomes. Structural and *in vitro* studies implicate Spt4 in transcription through nucleosomes, yet the *in vivo* function of Spt4 is unclear. Here we assessed the precise position of Spt4 during transcription and the consequences of loss of Spt4 on RNA polymerase II (RNAPII) dynamics and nucleosome positioning in *Saccharomyces cerevisiae*. In the absence of Spt4, the spacing between gene-body nucleosomes increases and RNAPII accumulates upstream of the nucleosomal dyad, most dramatically at nucleosome +2. Spt4 associates with elongating RNAPII early in transcription and its association dynamically changes depending on nucleosome positions. Together, our data show that Spt4 regulates early elongation dynamics, participates in co-transcriptional nucleosome positioning, and promotes RNAPII movement through the gene-body nucleosomes, especially the +2 nucleosome.

## Introduction

In eukaryotes, nucleosomes limit access to DNA and thus act as intrinsic barriers to DNA-dependent processes including RNA polymerase II (RNAPII) transcription (Kornberg, 1974; Zhou et al., 2019). Transcription requires the sequential breaking of interactions between nucleosomal DNA and histones, and reassembling the nucleosome after RNAPII has passed (Kujirai and Kurumizaka, 2020). A wide range of factors are implicated in assisting this route of RNAPII through nucleosomes but how they function in the cell is not yet clear (Clapier et al., 2017; Ehara et al., 2019; Farnung et al., 2018; Gurova et al., 2018; Venkatesh and Workman, 2015). One such factor is the DRB-sensitivity inducing factor (DSIF) complex in metazoans, also known as the Spt4/5 complex in yeasts, required for efficient transcription on chromatin (Crickard et al., 2017; Ehara et al., 2019; Kujirai and Kurumizaka, 2020; Vos et al., 2020). Spt4 is one of the most highly conserved transcription elongation factors (TEFs) in archaea and eukaryotes and its partner Spt5 is conserved in all three kingdoms, known as NusG in prokaryotes (Hartzog and Fu, 2013; Ponting, 2002). Spt4 and Spt5 are encoded by genes originally isolated as suppressors of the loss of gene expression as a result of the insertion of the budding yeast transposon, Ty, into a reporter gene (Winston *et al*., 1984), hence named *SPT* (Suppressor of Ty). Early experiments linked many of the *SPT* genes to transcription and chromatin, including *SPT6* (Swanson and Winston, 1992), *SPT16* (Malone et al., 1991), *SPT11*, and *SPT12* (Clark-Adams et al., 1988). Structural studies demonstrated that the Spt4/5 complex locates on top of the RNAPII active cleft and in between the upcoming nucleosome and RNAPII (Ehara et al., 2019; Farnung et al., 2018), *in vitro* transcription assays revealed that the Spt4/5 complex reduces RNAPII stalling during transcription through a nucleosome (Crickard et al., 2017; Ehara et al., 2019), and single-molecule experiments showed that the Spt4/5 complex differentially interacts with different RNAPII-nucleosome intermediates formed during transcription through the nucleosome (Crickard et al., 2017). Despite these studies implicating Spt4/5 in transcription regulation in the context of chromatin, the *in vivo* functions of Spt4/5 remain poorly understood. Furthermore, most studies focus on Spt4/5 as a complex which makes it hard to interpret the exact function of Spt4 and Spt5 as individual TEFs (Decker, 2020).

We used a combination of high-throughput sequencing and mathematical modelling approaches to investigate the exact function of Spt4 in transcription in the cell. Using NET-seq in the *spt4* knock-out (*spt4Δ*) cells to map the position of engaged RNAPII with base-pair resolution, mathematical modelling to predict the role of Spt4 in RNAPII dynamics, TEF-seq to map the precise position of Spt4 and Spt5 on RNAPII, and MNase-seq in *spt4Δ* cells, to investigate the impact of Spt4 on nucleosome positioning, we showed that the primary function of Spt4 is in early elongation. The association between Spt4 and RNAPII dynamically changes as RNAPII transitions through nucleosomes and gene-body nucleosome positions (from the +2 nucleosome onwards) are altered in *spt4Δ* cells. In both *spt4Δ* and cells depleted of Spt4 in real-time, RNAPII accumulates upstream of nucleosome dyads, especially at the +2 nucleosome. Overall, these findings support Spt4 promoting RNAPII movement through nucleosomes, especially in early transcription, and regulating co-transcriptional nucleosome positioning.

## Results

### In the absence of Spt4, RNAPII accumulates at the 5’end of genes

As Spt4 is an elongation factor, we asked whether Spt4 influences the genome-wide distribution of RNAPII using native elongating transcript sequencing (NET-seq). NET-seq maps the position of all forms of RNAPII with RNA in its active site, including paused or backtracked enzymes (Churchman and Weissman, 2011) (**Figure S1A**). Spike-in normalised NET-seq was performed in WT and *spt4* knock-out (*spt4Δ*) cells, also expressing FLAG-tagged RNAPII in biological duplicates. To remove the background signal, samples without FLAG-tag (no tag control) were processed in parallel to the tagged samples (see Methods). NET-seq repeats were reproducible and the NET-seq data presented here were consistent with the previously published NET-seq data (Fischl et al., 2017) (**Figure S1B**).

In WT cells, the NET-seq signal is relatively high within the first 500 nt from the transcription start site (TSS), then it drops while transcribing over the gene body, and peaks again upstream of the polyadenylation site (PAS) (Churchman and Weissman, 2011; Fischl et al., 2017) (**Figure 1A-C and S1C**). In *spt4Δ* cells, individual gene plots, heatmaps, and metagene plots showed that the density of RNAPII significantly increased over genes (**Figure 1A-D and S1C, D**). Importantly, the most apparent change in the distribution of RNAPII was within the first 200 nt from the TSS regardless of gene length (**Figure 1B, C and S1C, D**), suggesting that Spt4 regulates the distribution of RNAPII early in transcription.

**Figure 1).**
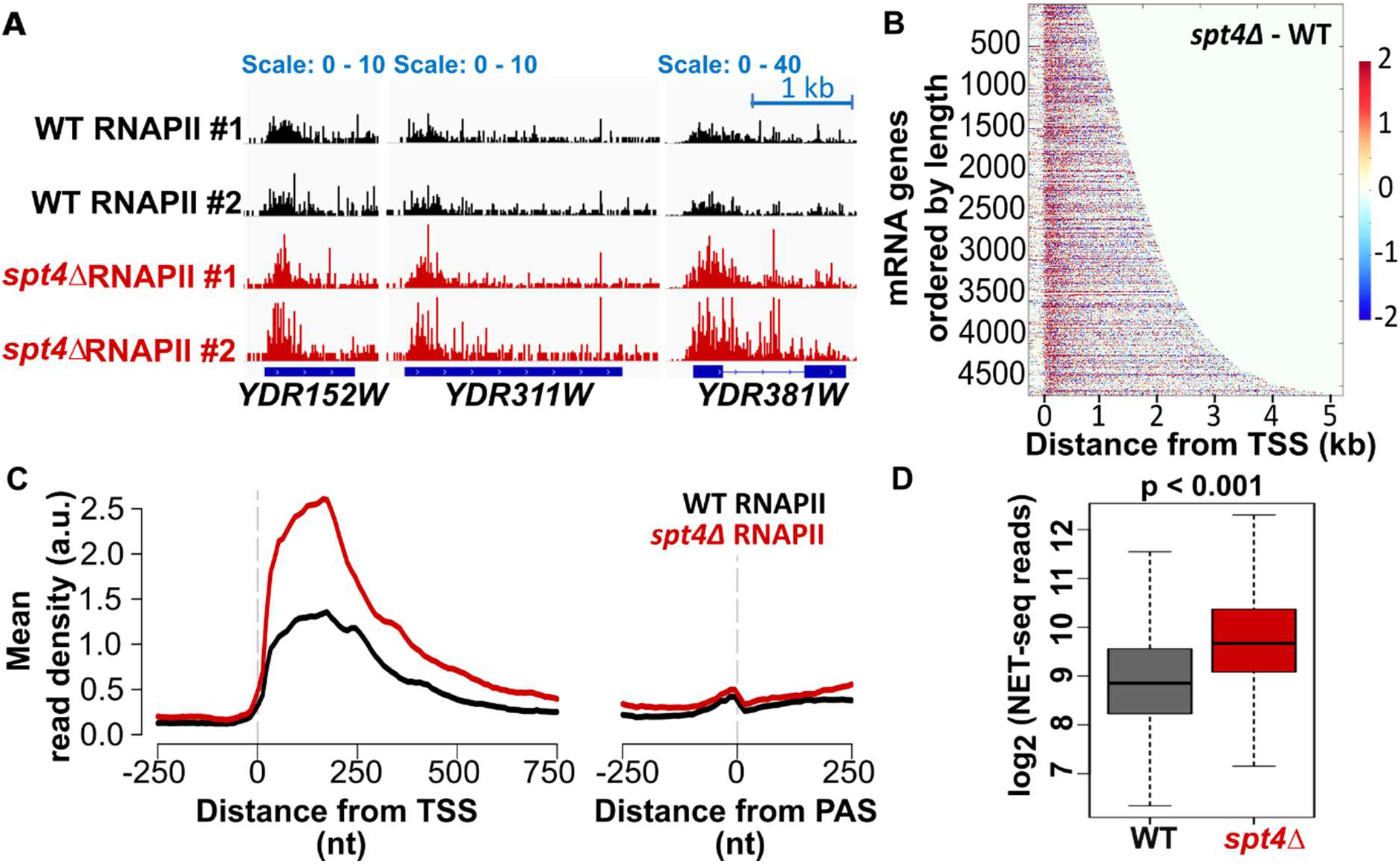
In the absence of Spt4, RNAPII accumulates at the 5’end of genes. **A)** WT and *spt4Δ* NET-seq signals of example genes transcribed from the positive strand: *YDR152W, YDR311W*, and *YDR381W* in two biological replicates. The dark blue boxes indicate the transcribed region of the genes (from TSS to PAS), the blue line indicates the intronic region in *YDR381W*. **B)** Heatmaps of the difference between the *spt4Δ* and WT NET-seq signal (*spt4Δ* - WT). Each row indicates a protein-coding gene (n=4610), ranked by gene length. The colour code reflects the changes in the RNAPII signal for each nucleotide position from TSS-250 nt to TSS+4750 nt (x-axis) as shown by the colour bar. **C)** Metagene plots of NET-seq reads in WT (black) and *spt4Δ* (red) aligned at the TSS or PAS. **D)** Boxplots of the NET-seq reads in WT (grey) and *spt4Δ* (red) cells on log_2_ scale. The reads were counted from TSS to PAS-250 nt for protein-coding genes after filtering low read genes out (see Methods). N=4610, p <0.001, two-tailed, paired Student’s t-test.

### Mathematical modelling supports defects in early transcription elongation in *spt4Δ* cells

To address potential mechanisms leading to the higher RNAPII signal in *spt4Δ* compared to WT cells, we developed a mathematical model designed to simulate the shape of the NET-seq profiles where a number of potential mechanisms occurring during transcription are ascribed relative numerical values (Brown, 2019) (**Figure S2**). This enabled us to relate the changes in the profiles of *spt4Δ* compared to WT cells to underlying transcription dynamics. The model computationally simulated RNAPII dynamics by considering initiation, elongation, occlusion of RNAPII by a downstream RNAPII, collision of RNAPIIs, stalling, backtracking, resolution of collision/backtracking/stalling events, and early termination. In contrasts to previous approaches (Azofeifa and Dowell, 2017; Fischer et al., 2020; Tufegdžić Vidaković et al., 2020), we set two distinct windows of transcription in which stalling and backtracking parameters can be different (**Figure 2A**), motivated by the notion that RNAPII is subject to distinct regulation in the early and late stages of transcription (Peck et al., 2019). The WT RNAPII position provided by experimental NET-seq data at single nucleotide resolution was fitted to the shape of the transcription profile simulated by the model. Modelling suggested that there are six key metrics that can be inferred from the shape of the WT RNAPII distribution: 1) ratio of the rate of initiation compared to elongation, 2) ratio of RNAPII moving compared to stalled or backtracked (moving ratio) in window 1, 3) the size of window 1, 4) the mean location of early termination, 5) the moving ratio in window 2, and 6) the processivity of RNAPII (% of initiating polymerase reaching 1000 nt) (**Figure 2A**). To test the extent of the change in each metric, the parameter values were obtained for each gene in WT and *spt4Δ* cells and the two conditions were quantitatively compared (**Figure 2B**). Three parameter values showed significant and marked differences in *spt4Δ* data compared to the WT. The increase in the initiation to elongation ratio suggested either a defect in elongation or an increased initiation frequency in the profiles from cells lacking Spt4. An overall defect in elongation is supported by a reduced proportion of moving polymerase in window 1 and a reduction in the processivity of polymerase in *spt4Δ* cells compared to WT (**Figure 2B**).

**Figure 2).**
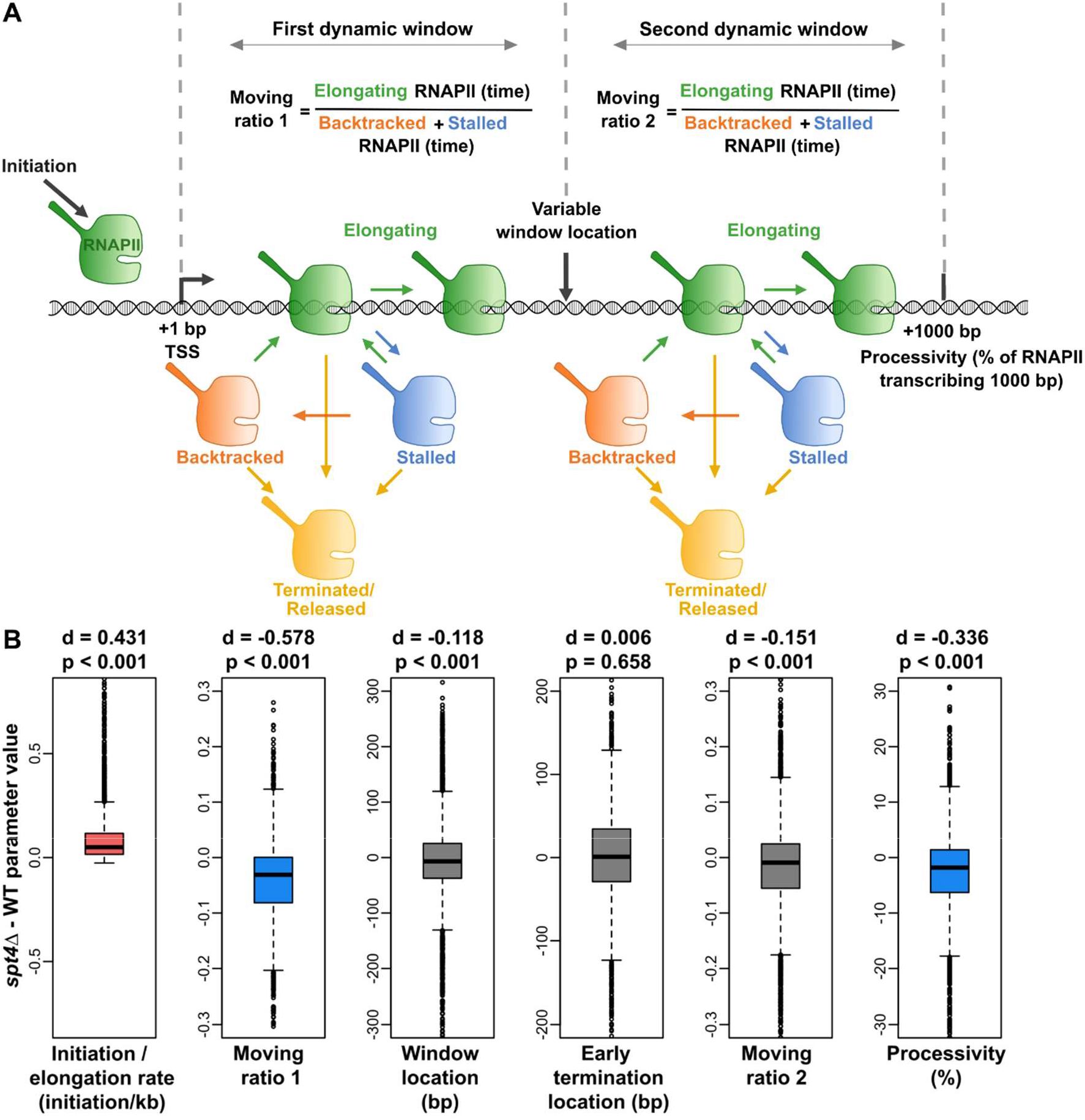
Mathematical modelling supports defects in early transcription elongation in *spt4Δ* cells. **A)** Schematic of the mathematical model. Model describes RNAPII transcription reaching 1000 bp with initiation rate (initiation per min), elongation rate (kb per min), stalling, backtracking, termination events (determined by Poisson distribution), variable window location (bp). Moving ratio 1 and 2 describes number of RNAPII elongating compared to backtracked or stalled RNAPII within respective transcription window. Processivity indicates % of RNAPII reaching 1000 bp. **B)** Metrics were obtained for each gene in WT and *spt4Δ*, and the two conditions were quantitatively compared. The significance of the changes was reported by calculating p-values and the magnitude of the changes was reported by calculating Cohen’s d (Cohen, 1988). Cohen’s d is computed by taking the mean difference between the WT and *spt4Δ* metric value divided by the standard deviation of the differences. The value of Cohen’s d gives a measure of the effect size of the change such that the values between 0.2 to 0.5 indicate small changes, between 0.5 and 0.8 indicate medium changes, and > 0.8 large changes (Cohen, 1988). Positive and negative values indicate a relative increase or decrease in the given metric, respectively. The red and blue boxplots indicate significant and marked increase and decrease, respectively, in the *spt4Δ* metric values compared to WT cells.

### The primary defect in *spt4Δ* cells is early transcription elongation

Modelling suggests that the movement of RNAPII in the early stages of transcription is the main defect in *spt4Δ* cells but does not allow us to distinguish whether this is a result of a defect at initiation or early elongation or both. To examine and validate the predictions of the model, we used three approaches: 1) we investigated the levels of the pre-initiation complex (PIC) at promoters as a proxy for transcription initiation frequency, 2) we compared elongation competent RNAPII with levels of all engaged RNAPII, and 3) we mapped RNAPII upon rapid depletion of Spt4 to detect the immediate changes in the distribution of RNAPII. Our data support a primary function for Spt4 in early transcription elongation, rather than initiation.

Sua7 (TFIIB) is a subunit of the pre-initiation complex (PIC) that is required for RNAPII recruitment to promoters (Sainsbury et al., 2015) and the amount of chromatin-bound Sua7 reflect the changes in transcription initiation levels (Doris et al., 2018). Therefore, we reasoned that if the transcription initiation rate was higher in *spt4Δ* cells, levels of Sua7 at promoters should also be higher compared to WT cells. To test this, we performed spike-in normalised ChIP-seq for Sua7 in WT and *spt4Δ* cells in biological duplicates and detected the Sua7 binding sites by peak-calling using MACS2 (Zhang et al., 2008). Sua7 ChIP-seq data was reproducible between the replicates (**Figure S3A, B**), and consistent with previous studies, genome-wide Sua7 mapped around the TSS (**Figure S3A**) (Doris et al., 2018). Visual inspection of individual genes and metagene plots revealed similar Sua7 occupancy in WT and *spt4Δ* cells (**Figure S3A and 3A**). Differential enrichment analysis of the Sua7 signal indicated no change in the level of the PIC in the absence of Spt4 (97.4%; 3148/3269) (**Figure 3B**). Consequently, these data do not support a change at initiation frequency in *spt4Δ* cell, in line with the *in vitro* studies suggesting the human counterpart of the Spt4/5 complex (DSIF) has no effect on the transcription initiation (Zhu et al., 2007).

**Figure 3).**
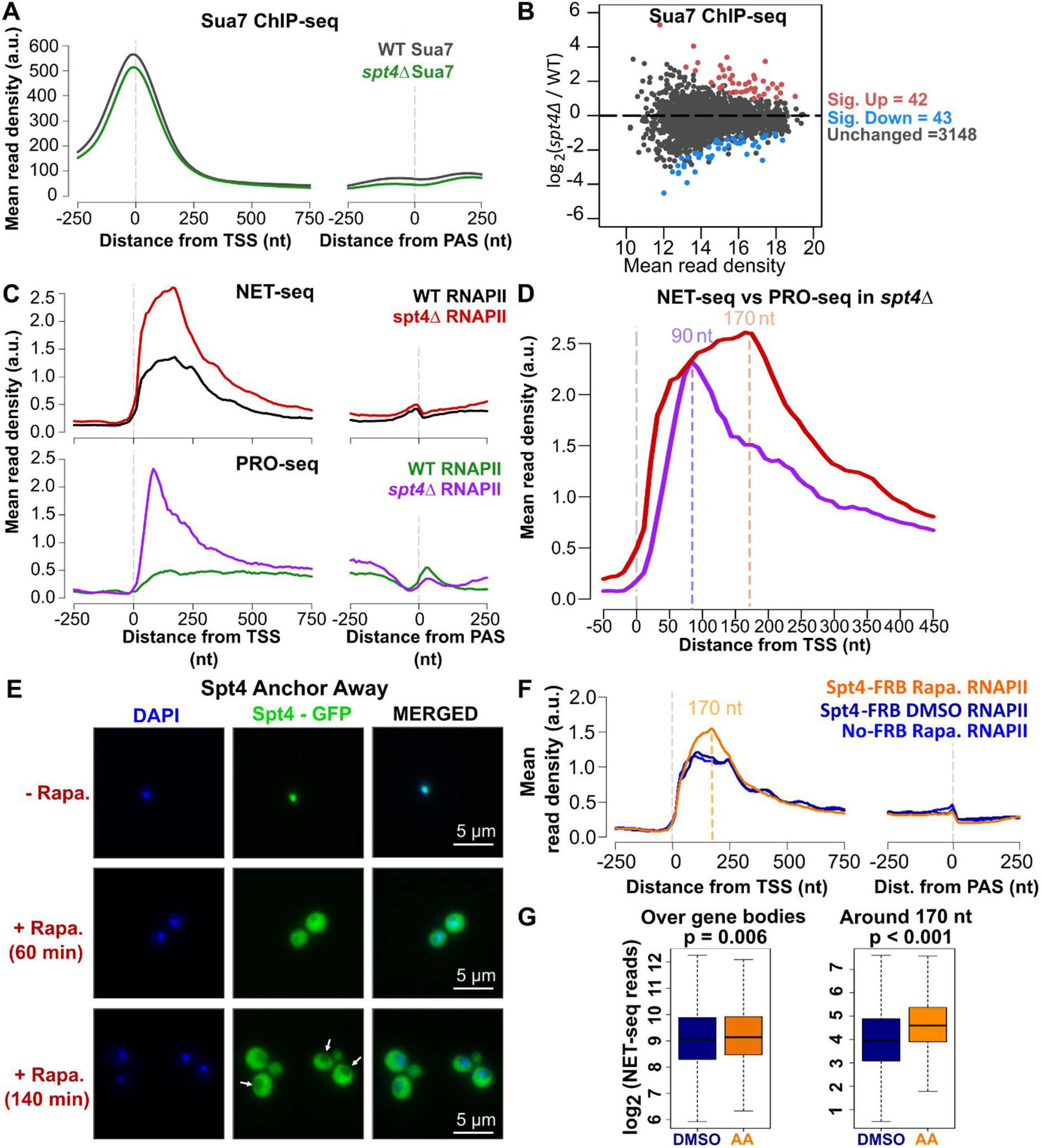
The primary defect in *spt4Δ* cells is early transcription elongation. **A)** Metagene plots of Sua7 ChIP-seq reads in WT (black) and *spt4Δ* (green) aligned at the TSS or PAS for protein coding genes (n=3233). **B)** Differential enrichment analysis of Sua7 in WT and *spt4Δ*. DEseq2 applied to the read counts around the TSS (TSS-100 to TSS+100 nt) for the two replicates of each data. Significantly enriched and depleted genes indicated in red and blue, respectively (p-adjusted <0.05). **C)** Metagene plots of NET-seq reads in WT (black) and *spt4Δ* (red) aligned at the TSS or PAS (the top panel), and metagene plots of published PRO-seq reads in WT (green) and *spt4Δ* (purple) aligned at the TSS or PAS (the bottom panel). PRO-seq data was taken from GEO:GSE76142 (Booth et al.,2016). **D)** Metagene plots of *spt4Δ* NET-seq (red) and *spt4Δ* PRO-seq (purple) reads aligned at the TSS, the same data as in C. Dashed lines indicate the highest PRO-seq (90 nt, purple) and NET-seq reads (170 nt, pink). **E)** IF images for Spt4-FRB samples at time points 0, 60, and 140 min after rapamycin addition. DAPI staining indicates nucleus, GFP is expressed with Spt4 (Spt4-FRB-GFP). **F)** Metagene plots of NET-seq reads in DMSO control (navy), rapamycin-treated Spt4-FRB (orange) and No-FRB cells (blue) aligned at the TSS or PAS. Dashed line indicates the highest NET-seq read upon Spt4 anchor away (170 nt, orange). **G)** Boxplots of the NET-seq reads in DMSO control (DMSO; navy) and rapamycin-treated Spt4-FRB (AA; orange) cells on log_2_ scale. Reads were counted for over gene bodies (TSS to PAS-250 nt) and at 170 +/- 10 nt from the TSS for protein-coding genes after filtering for low read genes (see Methods). N=4560, p=0.006 and p<0.001, respectively, two-tailed, paired Student’s t-test.

Another prediction of the model is a decreased moving ratio in early elongation (moving ratio 1), possibly due to more frequent, or less efficiently resolved, stalling and backtracking events. To distinguish between elongation competent and stalled/backtracked RNAPII in *spt4Δ* cells, we utilised precision-run-on sequencing (PRO-seq) profiles (Booth et al., 2016) and compared these to our NET-seq profiles. PRO-seq allows the mapping of RNAPII that is competent to elongate during the metabolic labelling period, while NET-seq captures all forms of RNAPII, including elongating, backtracked, and stalled. Therefore, if RNAPII is captured by NET-seq, but not PRO-seq, it would indicate a non-elongating but still RNA-engaged RNAPII (for example, backtracked) at a given position (**Figure 3C, D**). More stalled or backtracked RNAPII was observed in *spt4Δ* cells, particularly between ∼90 nt and ∼170 nt from the TSS (**Figure 3C, D**). This suggests that, in the absence of Spt4, RNAPII transcribes with short term pauses to around 90 nt from the TSS, whereas between 90 and 170 nt, more of the RNAPII is stalled or backtracked, leading to the decreased moving ratio in window 1, supporting an early elongation defect.

### Rapid depletion of Spt4 leads to the accumulation of RNAPII around 170 nt into gene bodies

We monitored the effect of real-time loss of the Spt4 protein from the nucleus using the anchor-away system (AA) (Haruki et al., 2008) on the distribution of RNAPII. The AA allows the conditional removal of a target protein from the nucleus upon rapamycin addition. The efficient depletion of Spt4 was verified by immunofluorescence microscopy (**Figure 3E**) and quantified by the effect on growth rate (**Figure S3C**) as well as by ChIP-qPCR (**Figure S3D**). NET-seq was performed in biological duplicate in Spt4 anchor away cells (Spt4-AA). DMSO treated Spt4-FRB-GPF and rapamycin-treated No-FRB cells were included as controls. Spt4-AA NET-seq repeats and control experiments were reproducible (**Figure S3E, G**). Intriguingly, the real-time depletion of Spt4 has a small effect on the distribution of RNAPII across gene bodies but led to the most notable and significant changes in the RNAPII profile around 170 nt from the TSS (**Figure 3F, G**). This complements the *spt4Δ* NET-seq results and demonstrates that the change in the distribution of RNAPII in the absence of Spt4 first manifests itself around 170 nt downstream from the TSS.

### The Spt4/5 complex travels with RNAPII

Next, we assessed where Spt4 and Spt5 associate with RNAPII to examine whether the Spt4/5 complex is particularly enriched on RNAPII where the effect on transcription elongation is most marked (at the 5’ end of genes). To this end, we investigated their genome wide positions on RNAPII during transcription using transcription elongation factor (TEF) associated nascent elongating transcript sequencing (TEF-seq) (Fischl et al., 2017). TEF-seq is a variation of NET-seq (**Figure S4A**), giving single nucleotide-resolution mapping of the position of the RNAPII-associated TEFs (Fischl et al., 2017).

Spt4 or Spt5 was FLAG-tagged for immunoprecipitation of the factor-associated transcription complex. Samples without FLAG-tag (no tag control) were included to remove the background signal as described for the NET-seq. Spike-in normalised Spt4 and Spt5 TEF-seq were performed in duplicate, and the results were reproducible (**Figure 4A and S4B**). The Spt4 and Spt5 signals are similar over gene bodies, consistent with Spt4 and Spt5 forming a highly stable complex (Hartzog et al., 1998) (**Figure 4A, B**). The Spt4 and Spt5 profiles match the RNAPII (NET-seq) profile, suggesting engagement of these factors with RNAPII throughout transcription (**Figure 4A, B**) agreeing with the low resolution chromatin distribution provided by previous ChIP-based studies (Mayer et al., 2010). Importantly, here, TEF-seq allows the detection of co-transcriptional and native interactions between TEFs and RNAPII at single nucleotide resolution. At the 5’-end of the genes, Spt4 and Spt5 signals rise as early as the RNAPII signal (**Figure 4B**) implying that Spt4/5 engages with RNAPII at the early stage of transcription, consistent with the structural and *in vitro* findings showing that Spt4/5 replace transcription initiation factors and is recruited to the transcription elongation complex once 20 nt RNA has been transcribed by RNAPII (Bernecky et al., 2017; Grohmann et al., 2011; Rosen et al., 2020). Over the gene bodies, Spt4/5 remain associated with elongating RNAPII (**Figure 4B**). At the 3’-end of the genes, Spt4/5 signals drop to the background levels around 100 nt before the polyadenylation site (PAS) (**Figure 4B and S4C**). This region coincides with the 3’end peak of RNAPII where pausing and reduced speed of RNAPII is predicted to be important for the transition from elongation to termination complex (Mischo and Proudfoot, 2013). ChIP-seq studies in *S*.*pombe* suggest that Spt5 dissociates from the RNAPII during the transition from the elongation to the termination complex (Kecman et al., 2018; Parua et al., 2018). The drop in the Spt5/4 signal is likely to reflect a similar transition mechanism in budding yeast, allowing the binding of termination factors. Overall, the data indicate Spt4/5 joining RNAPII right after initiation, travelling with elongating RNAPII, and dissociating from RNAPII about 100 nt upstream of the PAS.

**Figure 4).**
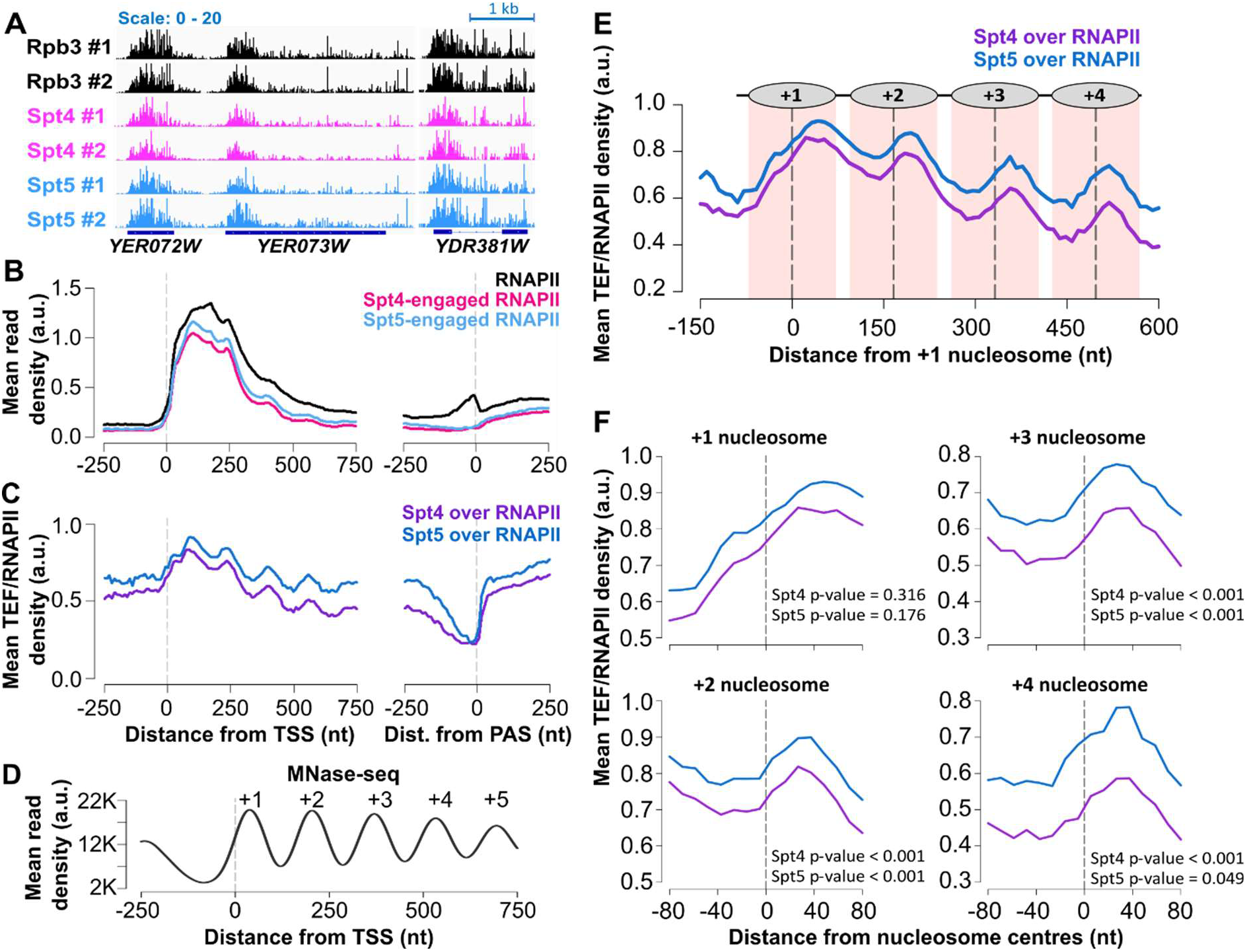
Spt4/5 travel with RNAPII and oscillate on and off RNAPII based on the nucleosome positions. **A)** NET-seq (RNAPII) and TEF-seq (Spt4 and Spt5) reads of example genes transcribed from the positive strand: *YER072W, YER073W*, and *YDR381W* in two biological replicates. The dark blue boxes indicate the transcribed region of the genes (from TSS to PAS), the blue line indicates the intronic region in *YDR381W*. **B)** Metagene plots of NET-seq (RNAPII; black), and TEF-seq (Spt4; pink, Spt5; light blue) reads aligned at the TSS or PAS. **C)** Metagene plots of Spt4 over RNAPII (purple) and Spt5 over RNAPII (dark blue) data aligned at the TSS or PAS. Spt4 over RNAPII was plotted by dividing Spt4-engaged RNAPII signal (TEF-seq) by RNAPII signal (NET-seq). The same is applied to Spt5 TEF-seq data. **D)** Metagene plots of MNase-seq reads in WT cells aligned at the TSS. The dashed line in grey indicates the TSS. **E)** Metagene plots of Spt4 over RNAPII (purple) and Spt5 over RNAPII (dark blue) relative to the +1 nucleosome dyad. Dashed lines (black) through the peaks indicate the centres of the nucleosomes and the nucleosomal DNA (+/- 70 nt around the centre) is highlighted in light pink. Position of nucleosomes graphically shown above the metagene plot. **F)** Spt4 and Spt5 occupancies on RNAPII are shown around individual nucleosome dyads +1,+2,+3, and +4. The TEF/RNAPII values from upstream of the dyad (−60 to -10 nt from the dyad) and downstream of the dyad (+10 to +60 nt from the dyad) were compared for each gene. Significance of the change in the factor occupancies around the nucleosomes were tested by one tailed (condition: upstream signal < downstream signal), paired Student’s t-test.

Additionally, to test whether Spt4/5 were differentially enriched for specific groups of genes, we plotted RNAPII, Spt4, and Spt5 signal as heatmaps based on RNAPII occupancy level and performed a quantitative analysis of genome wide Spt4 and Spt5 occupancies on RNAPII. The Spt4 and Spt5 levels were proportional to the RNAPII levels at most genes (>99%) and thus Spt4/5 participates in transcription of nearly all mRNA genes (**Figure S4D-F**).

### Spt4/5 oscillate on and off RNAPII based on nucleosome positions

The TEF-seq signals for Spt4/5 come from the native RNA attached to RNAPII associated with Spt4/5. To demonstrate relative occupancies of Spt4 and Spt5 on RNAPII, we plotted the TEF-seq signal relative to the NET-seq signal. Interestingly, the association of Spt4/5 with RNAPII was not constant, but periodically changing (**Figure 4C**). As their periodicity resembles the frequency of nucleosome phasing, we compared the NET-seq normalised TEF-seq profiles with nucleosome positions, derived using micrococcal nuclease (MNase) digestion followed by DNA sequencing (MNase-seq) in WT cells. MNase-seq was performed in biological triplicates and analysed using DANPOS2 to compute estimates for the nucleosome protected reads and nucleosome dyad positions (Chen et al., 2013). In a typical MNase-seq profile, nucleosome depleted regions (NDRs) are detected at promoters, and nucleosomes are regularly arrayed in gene bodies relative to the transcription start sites (TSS) (Baldi et al., 2020) and our MNase-seq resulted in the expected and reproducible digestion pattern across the genome (**Figure 4D, S5A**).

Next, we re-plotted the NET-seq normalised TEF-seq profiles relative to the +1 nucleosome dyad to test if nucleosome positions correlate with the phasing patterns of Spt4/5. Remarkably, the oscillation pattern of the Spt4/5 on RNAPII was off-set with respect to nucleosome positions (**Figure 4E**). More detailed analysis was done by plotting the metagene profiles around the nucleosome dyads (+1 to +4), separately (**Figure 4F**). The Spt4 and Spt5 occupancies on RNAPII were not different around the +1 nucleosome (**Figure 4B, D**). However, on the gene-body nucleosomes (+2 to +4), the Spt4 and Spt5 occupancies on RNAPII were significantly lower at the upstream face of the nucleosome dyads compared to the downstream face (**Figure 4F**). As RNAPII transcribes through nucleosomes, RNAPII-nucleosome conformations change. *In vitro* studies from Crickard *et al*. documented that Spt4/5 stabilises the RNAPII-nucleosome intermediate after RNAPII passing the dyad (Crickard et al., 2017) and this is where we observe the higher levels of the Spt4/5 complex on RNAPII. Together, these results support Spt4/5 dynamically interacting with the transcription elongation complex during transcription and raises the question as to whether Spt4 and Spt5 have a direct and equivalent role in chromatin transcription.

### Spt5 and Spt4 have distinct impacts on transcription

As expected from factors in a complex, Spt4 and Spt5 show similar patterns of association with RNAPII across genes, including oscillations (**Figure 4A-C**). As studies suggested that Spt4 is important for the stability of Spt5 (Ding et al., 2010; Krasnopolsky et al., 2021), we also mapped the position of Spt5 during transcription in *spt4Δ* cells using TEF-seq (**Figure 5A, S5A, B**). The levels of Spt5 on RNAPII were reduced in the absence of Spt4 and interestingly, the oscillation of Spt5 on RNAPII were also lost, supporting a role for Spt4 in stabilising/recruiting Spt5 to polymerase and in the oscillations of the complex on RNAPII as it transcribes through nucleosomes (**Figure 5B-D**). Notably, as shown in WT cells, in *spt4Δ* cells, the Spt5 level was proportional to the RNAPII level at most genes (>99%), implying that Spt5 was not differentially recruited to genes in the absence of Spt4 (**Figure S5C, D**).

**Figure 5).**
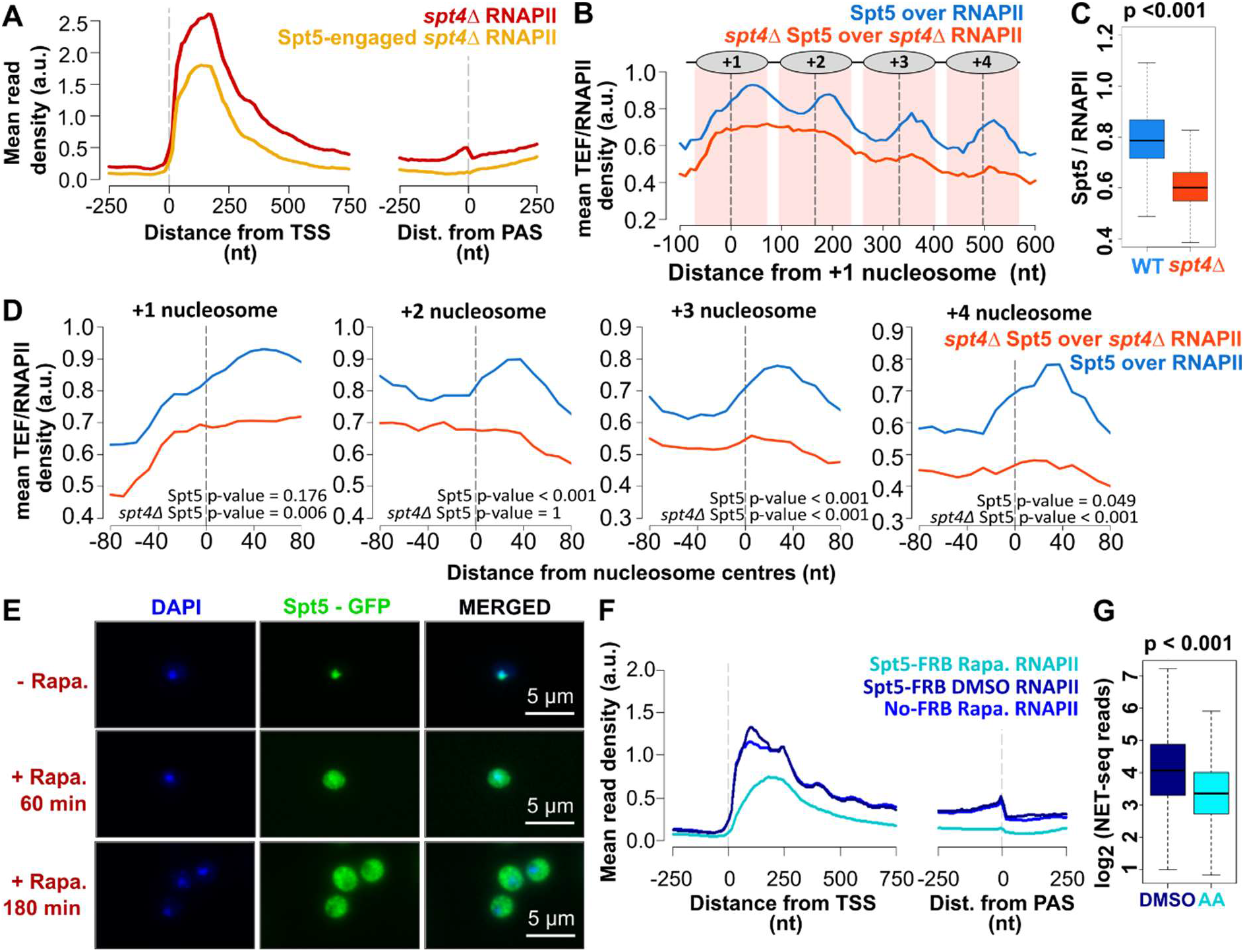
Spt5 and Spt4 have distinct impacts on transcription. **A)** Metagene plots of *spt4Δ* NET-seq (RNAPII; red), and *spt4Δ* Spt5 TEF-seq (yellow) reads aligned at the TSS or PAS. **B)** Metagene plots of Spt5 over RNAPII (blue) and *spt4Δ* Spt5 over *spt4Δ* RNAPII (orange) relative to the +1 nucleosome dyad. Plotted as described in Figure 4E. **C)** Boxplots of Spt5 over RNAPII (blue) and spt4Δ Spt5 over *spt4Δ* RNAPII (yellow) ratios for the protein-coding genes. The medians of the ratios (0.79 and 0.60, respectively) calculated by taking the reads from gene bodies (TSS to PAS-250 nt) of Spt5 TEF-seq and dividing by the reads from gene bodies of NET-seq both in WT and *spt4Δ*. (p-value < 0.001, Student’s t-test, paired, two-tailed). **D)** Spt5 occupancies on RNAPII in WT or *spt4Δ* cells are shown around individual nucleosome dyads +1,+2,+3, and +4. Plotted and tested as described in Figure 4F. **E)** IF images for Spt5-FRB samples at time points 0, 60, and 180 min after rapamycin addition. DAPI staining indicates nucleus, GFP is expressed with Spt5 (Spt5-FRB-GFP). **F)** Metagene plots of NET-seq reads in DMSO control (navy), rapamycin-treated Spt5-FRB (cyan) and No-FRB cells (blue) aligned at the TSS or PAS. **G)** Boxplots of the NET-seq reads in DMSO control (DMSO; navy) and rapamycin-treated Spt5-FRB (AA; cyan) cells on log_2_ scale. Reads were counted for over gene bodies (TSS to PAS-250 nt). P-value <0.001, respectively, two-tailed, paired Student’s t-test.

Next, to distinguish whether Spt4 and Spt5 have similar impact on transcription, using NET-seq, we examined the distribution of RNAPII over the genes upon Spt5 anchor away. The efficient depletion of Spt5 was verified by immunofluorescence microscopy (**Figure S5E**) and quantified by the effect on growth rate (**Figure S3C**) and ChIP-qPCR (**Figure S5E**). NET-seq was performed in biological duplicate in Spt5 anchor away cells (Spt5-AA). DMSO treated Spt5-FRB-GPF and rapamycin-treated No-FRB cells were included as controls. Spt5-AA NET-seq repeats and control experiments were reproducible (**Figure S5F, G**). Loss of Spt5 resulted in significant loss of NET-seq reads across the whole of the gene body, including at the polyadenylation site (**Figure 5F, G**). These results demonstrated that the NET-seq profiles upon Spt4 or Spt5 depletion were distinctly different (**Figure 3F and S3D**) consistent with an additional, essential function for Spt5 in transcription compared to Spt4 (Shetty et al., 2017). As Spt5 affects transcription so dramatically, we focussed only on Spt4 and asked whether Spt4 influences the organisation of nucleosomes with the aim of explaining its effect on RNAPII distribution.

### Spt4 influences nucleosome positioning

We have previously observed an oscillating pattern of Spt6 and Spt16 on RNAPII (Fischl et al., 2017) and interestingly, mutations in *spt6* and *spt16* have major effects on nucleosome positions (Doris et al., 2018; Feng et al., 2016). Therefore, next we asked whether Spt4 has an impact on nucleosome arrangement using MNase-seq in *spt4Δ* cells. Three replicates of MNase-seq in *spt4Δ* cells produced reproducible digestion patterns (**Figure S6A**).

Genome wide analysis was performed for protein coding genes (PCGs) longer than 600 nt and with well-defined nucleosome peaks (see Methods). Interestingly, in *spt4Δ* cells, although MNase digestion resulted in well-defined peaks across the genome (**Figure 6A, B and S6A**), nucleosomes were shifted towards the 3’-end of the genes compared to WT cells (**Figure 6B**). The differences between the nucleosome positions were more apparent at downstream nucleosomes. More detailed analysis was performed by calculating the position of the -1, +1, +2, +3, and +4 nucleosomes relative to the transcription start site (TSS) in the three replicates of the MNase-seq data. There was no difference in the median positions of the -1 and +1 nucleosomes in WT and *spt4Δ* cells (**Figure 6C and S6B**). Consistently, NDR length (defined as the distance between the dyads of the -1 and +1 nucleosome) in WT and *spt4Δ* cells were not different (**Figure S6C**). By contrast, the position of the +2, +3, and +4 nucleosome in *spt4Δ* cells were progressively shifted 3’ compared to WT cells (**Figure 6C and S6B**) suggesting increased nucleosome spacing (defined as the distance between the dyads of adjacent nucleosomes) in *spt4Δ* cells. Indeed, nucleosome spacing between nucleosome pairs (+1 to +2, +2 to +3, and +3 to +4) in *spt4Δ* were larger compared to WT cells (**Figure S6C**). Overall, the data support a role for Spt4 in positioning the gene-body nucleosomes from the +2 nucleosome, but no role in the positioning of the -1 and +1 nucleosomes. This suggests that Spt4 impacts nucleosome positioning during transcription elongation.

**Figure 6).**
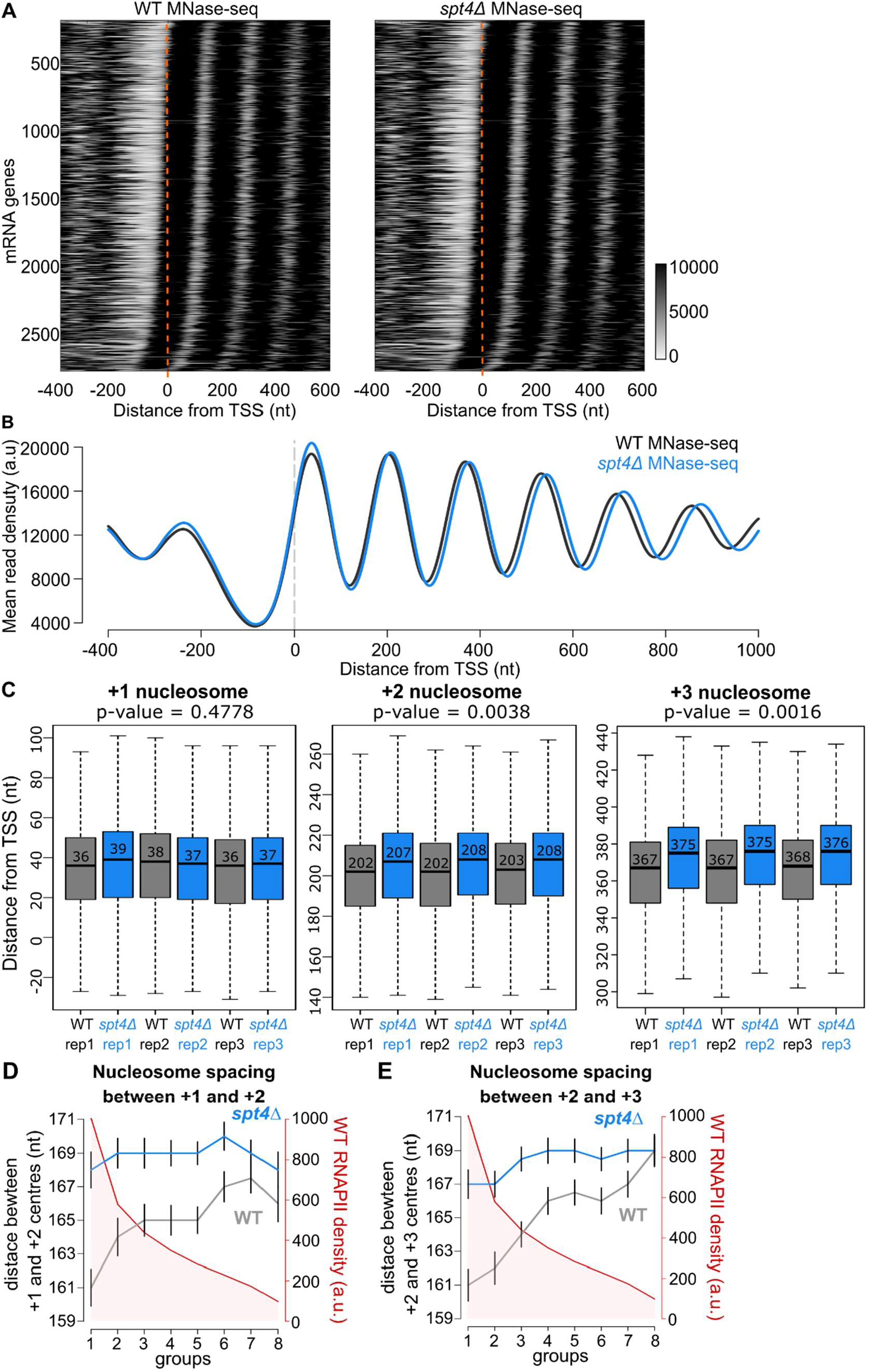
Spt4 influences nucleosome positioning. **A)** Heatmaps of MNase-seq reads in WT (left) and *spt4Δ* (right). PCGs ordered based on the position of +1 nucleosome in WT. Dashed line in orange indicates the TSS. **B)** Metagene plots of MNase-seq reads in WT (black) and *spt4Δ* cells (blue) aligned at the TSS. The dashed line in grey indicates the TSS. **C)** Box-plots of the distance of the +1, +2, and +3 nucleosomes from the TSS in three biological replicates of WT (black) and *spt4Δ* cells (blue). Numbers in the boxes indicate the median position of the given nucleosome. p-values were calculated by comparing the median position of the +1, +2, or +3 nucleosomes in WT and *spt4Δ* conditions obtained from each replicate (Student’s t-test, paired, two sided). **D)** Protein-coding genes were split into 8 groups based on WT NET-seq reads in the first 500 nt from the TSS (red, y-axis). The median NET-seq reads and the median nucleosome spacing between +1 and +2 nucleosomes in *spt4Δ* (blue) and WT (grey) were plotted for each group. The black bars around nucleosome spacing data points indicate one standard deviation. **E)** The median nucleosome spacing between +2 and +3 nucleosomes in *spt4Δ* (blue) and WT (grey) were plotted for each group as in D.

### Close nucleosome spacing at highly transcribed genes is dependent on Spt4

If Spt4 co-transcriptionally regulates nucleosome positioning, nucleosome spacing should be affected by the deletion of Spt4 to a greater extent in highly active genes compared to less active genes. To test this hypothesis, we investigated the correlation between nucleosome spacing and RNAPII densities (**Figure 6D, E**). PCGs were split into 8 groups based on their RNAPII density assessed by NET-seq reads. For each group, the median RNAPII density and the median nucleosome spacing between the +1 and +2, as well as between the +2 and +3 nucleosomes in WT and *spt4Δ* were plotted together. In WT cells, nucleosome spacing was shorter in highly expressed genes and it progressively increased for the genes having lower expression levels (**Figure 6D, E**). In *spt4Δ* cells, the distance between the nucleosomes was less variable, and larger than that of WT in all groups (**Figure 6D, E**). In other words, there was an overall increase in nucleosome spacing in *spt4Δ* cells compared to WT cells, and the increase was larger for highly expressed genes. In conclusion, the analysis shows that close nucleosome spacing observed in highly transcribed genes was lost in the absence of Spt4.

### The accumulation of RNAPII in the absence of Spt4 is associated with the position of the +2 nucleosome

The dynamic interaction of Spt4 with RNAPII based on nucleosome positions and the impact of Spt4 in gene-body nucleosome positions point to a role for Spt4 in chromatin transcription. Furthermore, the accumulation of RNAPII around 170 nt from the TSS in the absence of Spt4 brings about the possibility of a transcriptional barrier around this point. As *in vitro* studies have shown that the Spt4/5 complex does not help RNAPII progress over non-nucleosomal transcription barriers (Xu et al., 2020), but does aid RNAPII movement over nucleosomal barriers (Ehara et al., 2019; Farnung et al., 2020), we sought *in vivo* evidence for this by investigating the change in the distribution on RNAPII relative to nucleosome positions in the absence of Spt4.

Mapping the RNAPII density from normalised NET-seq reads to the position of the nucleosome dyads (+1 to +4) revealed the position of RNAPII accumulation at the upstream face of the +2 nucleosome *spt4Δ* cells, and to a lesser extent on the +3 and +4 nucleosomes (**Figure 7A, B**). Importantly, the same was also observed in Spt4-AA cells on the +2 nucleosome (**Figure 7C, D**), verifying the impact of the loss of Spt4 in the RNAPII distribution relative to nucleosomes in two different backgrounds. These results suggest that the main function of Spt4 is preventing RNAPII accumulation at the upstream face of nucleosomes, most specifically at the +2 nucleosome. As our modelling and experimental data pointed to increased stalling or backtracking during early elongation, we compared the NET-seq profile from *spt4Δ* cells and cells lacking Dst1 (TFIIS) (**Figure S7A, B**). Dst1 is a TEF that helps rescue backtracked RNAPII by triggering the cleavage activity of RNAPII (Zatreanu et al., 2019). Furthermore, loss of Dst1 is reported to lead to RNAPII accumulation around the nucleosome dyads (Churchman and Weissman, 2011). Interestingly, the RNAPII profiles in *spt4Δ* and *dst1Δ* are quite distinct, with Dst1 function focused on the dyad region of the +1 nucleosome (**Figure 7A, B**). This confirms that reads around the +1 nucleosome can be detected using NET-seq and supports a specific function for Spt4 in elongation at the +2 nucleosome. Taken together, we propose that the *in vivo* function of Spt4 involves helping RNAPII pass nucleosomal barriers downstream of the +1 nucleosome, especially at the +2 nucleosome (**Figure 7E**).

**Figure 7).**
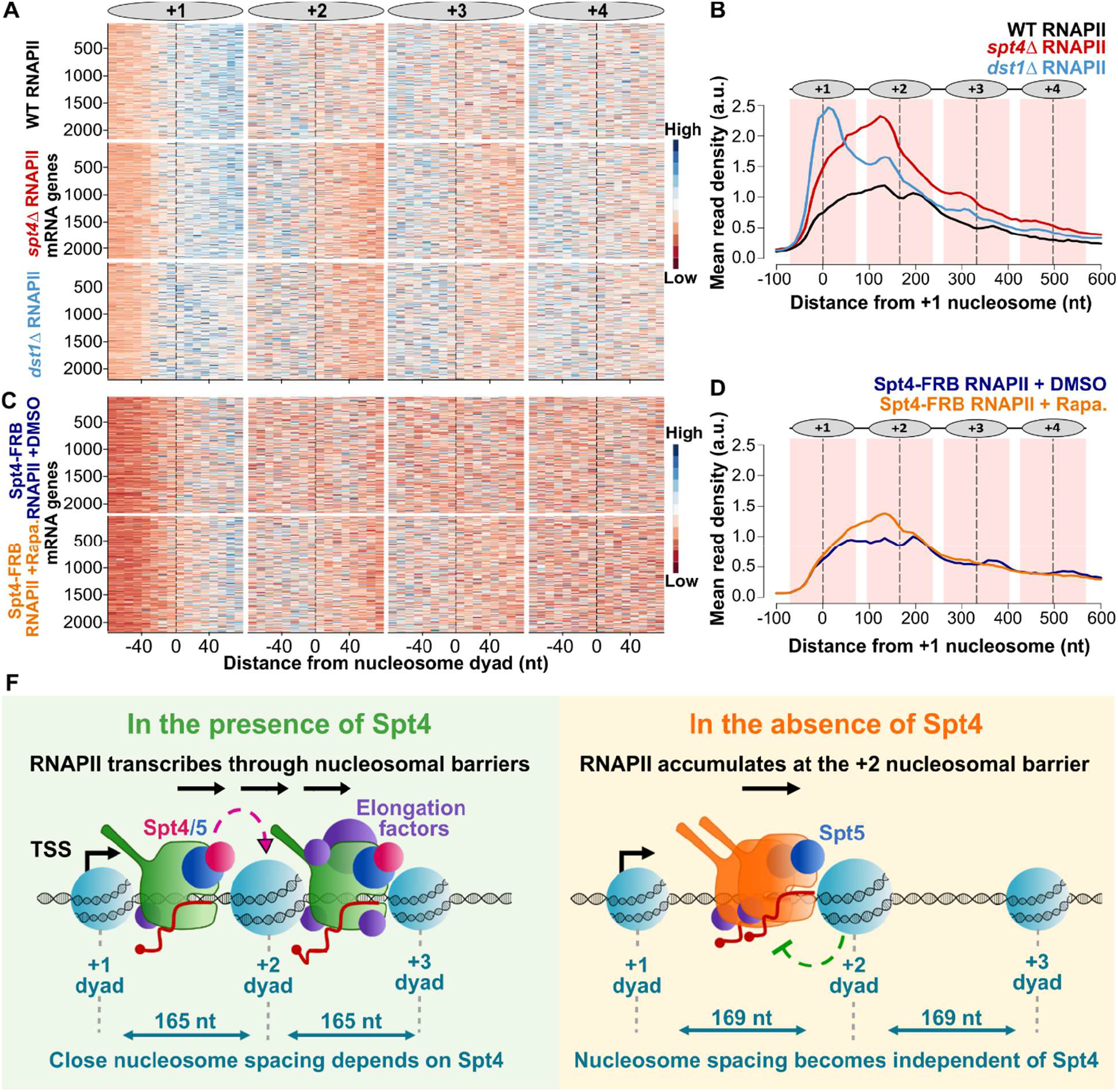
The accumulation of RNAPII in the absence of Spt4 is associated with the position of the +2 nucleosome. **A)** Heatmaps of WT (top), *spt4Δ* (middle), and *dst1Δ* (bottom) NET-seq profiles around the +1, +2, +3 and, +4 nucleosomes. Each row indicates a PCG (n=2212). RNAPII signal is shown in 10 nt bins around the indicated nucleosome dyads (−/+ 80 nt from the dyad; x-axis). The NET-seq reads were normalised to the mean and standard deviation of each gene to indicate the shape of the distribution of RNAPII regardless of the expression level differences between the genes. **B)** Metagene plots of WT (black), *spt4Δ* (red), and *dst1Δ* (blue) NET-seq profiles relative to the +1 nucleosome dyad. Dashed lines (black) through the peaks indicate the centres of the nucleosomes and the nucleosomal DNA (+/- 70 nt around the centre) is highlighted in light pink. Position of nucleosomes graphically shown above the metagene plot. **C)** Heatmaps of DMSO-treated Spt4-FRB (top) and rapamycin-treated Spt4-FRB (bottom) NET-seq profiles. Plotted as described in A. **D)** Metagene profiles of DMSO-treated (navy) and rapamycin-treated (orange) Spt4-FRB NET-seq profiles. Plotted as described in B. **F)** Model for Spt4 function.

## Discussion

Although structural and *in vitro* studies implicated Spt4/5 in RNAPII movement through nucleosomes, their precise role in transcription in the cell is poorly defined. Here, we reveal that Spt4/5 associates with RNAPII early in transcription and travels with elongating RNAPII over the gene bodies. As RNAPII transcribes over nucleosomes, the association of Spt4/5 with RNAPII oscillates, being higher at the downstream face of the dyad. Although Spt4 and Spt5 show similar distributions on RNAPII, Spt4 and Spt5 have different effects on RNAPII density over genes. Spt4 leads to accumulation of RNAPII at the 5’-end of genes, particularly at the upstream face of the +2 nucleosome, and to a lesser extent at the upstream face of the +3 and +4 nucleosomes. Interestingly, the accumulation of RNAPII on nucleosomes occurs at positions where levels of Spt4 are lowest. Finally, we show that in the absence of Spt4, the positions of the gene-body nucleosomes (+2 and beyond) are shifted downstream. Together, our data point to a primary role for Spt4 in regulating the movement of RNAPII through the +2 nucleosomal barrier.

Could Spt4 use the same mechanism to influence the nucleosome-related oscillations on RNAPII, the efficient movement of RNAPII through the +2 nucleosomal barrier, and nucleosome spacing? We considered two possibilities: an interaction with histones and/or with the nucleosomal DNA.

Like the Spt4/5 complex, the histone chaperones Spt6 and Spt16 also oscillate, out of phase, on and off RNAPII, reflecting their dynamic interactions with different histones during transcription (Fischl et al., 2017). This raises the possibility of distinct affinities by these different TEFs for specific conformations of histones with elongating RNAPII, leading to the oscillations. Like histone chaperones, Spt5 bears an acidic domain that is predicted to interact with H2A/H2B during transcription (Ehara et al., 2019; Farnung et al., 2020). Spt4 does not have charged domains, but the affinity of Spt4 for nucleosomes could change indirectly through Spt5. The second possibility is binding of the Spt4/5 complex to nucleosomal DNA as it peels off from the nucleosome while RNAPII is moving forward. Indeed, Spt5 interacts with free DNA *in vitro* suggesting such dynamics between the Spt4/5 and DNA is also possible (Crickard et al., 2016). Either through an interaction with histones or nucleosomal DNA (or both), our model supports a function for Spt4 facilitating RNAPII movement on the nucleosomal barriers and aligns well with an *in vitro* model suggesting that together with FACT or Chd1, Spt4/5 contributes to effective RNAPII transcription through a nucleosome (Farnung et al., 2020).

This function of Spt4 would also explain why RNAPII accumulates at the upstream face of the nucleosomes. As *spt4Δ* cells are viable, RNAPII appears to pass nucleosomal barriers by redundant mechanisms but, these might be less effective at the +2 nucleosome, which was also recently recognised as an important barrier in stress (Badjatia et al., 2021). Additionally, dynamic changes in the composition of the transcription elongation complex as RNAPII transcribes along the genes could explain why the most notable effect of the loss of Spt4 is at the +2 nucleosome. Factors such as the Paf1 complex (Paf1C), are recruited to RNAPII around the +2 nucleosome and its level on RNAPII progressively increases towards the 3’-end of genes (Fischl et al., 2017). In the absence of Paf1, RNAPII accumulates at the downstream face of the +2 nucleosome (**Figure S7C, D**). Paf1C is a TEF complex generally associated with productive elongation as it takes part in co-transcriptional histone PTMs (Van Oss et al., 2017) and increases the processivity of RNAPII *in vitro* (Vos et al., 2020). Therefore, the movement of RNAPII through the upstream face of the +2 nucleosome might rely more on the function of Spt4. Around the +3 and +4 nucleosomes, Spt4 still contributes to transcription, possibly providing allosteric interactions. This could also explain the synthetic lethality in the double mutants of *spt4* and genes encoding the five Paf1C components (Squazzo et al., 2002). Alternatively, the reason why Spt4 is most crucial for passing the +2 nucleosome might be related to specific histone post-translational modifications (PTMs). The role of histone PTMs in overcoming nucleosome barriers remains unknown and future studies will be needed to investigate this.

The negative correlation between nucleosome spacing and the RNAPII density on genes observed here and by others (Baldi et al., 2018; Ocampo et al., 2016) could result from high levels of transcription causing either removal or re-positioning of nucleosomes to allow RNAPII passage (Singh et al., 2021). This would, in turn, lead to a delay in restoration of normal spacing, which is an energy requiring process involving remodellers such as Isw1 and Chd1 (Gkikopoulos et al., 2011; Kent et al., 2001; Morillon et al., 2003; Ocampo et al., 2019). Here, our model would also explain the increased nucleosome spacing observed in the absence of Spt4. If Spt4 helps RNAPII pass nucleosomal barriers, inefficient removal or re-positioning of nucleosomes would eliminate the need for restoration of nucleosome positioning which would also explain the observations suggesting opposing roles for Isw1 and Spt4 in transcription through chromatin (Morillon et al., 2003).

Finally, we considered a role for Spt4 in transcription itself. Our mathematical model predicts and others report that, in *SPT4* mutants, RNAPII shows an elongation defect and is less processive (Booth et al., 2016; Hartzog and Fu, 2013; Mason and Struhl, 2005). This must be balanced by a reduction in transcript turnover rates (Brown et al., 2018), as overall levels of transcripts do not change in *spt4Δ* cells (Booth et al., 2016). The increased NET-seq signal would also be consistent with an elongation defect in *spt4Δ* cells. Is an elongation defect linked to the nucleosome spacing defect, which is similar to a pattern that is normally observed in lowly expressed genes or upon RNAPII depletion (Singh et al., 2021; Weiner et al., 2010), or to accumulation of RNAPII on nucleosomes? Work with other mutants suggests no simple relationship between accumulation of RNAPII upstream of nucleosomes, reduced RNAPII processivity, and increased nucleosome spacing. For example, *hpr1Δ* mutants have less processive RNAPII (Mason and Struhl, 2005) and *dst1Δ* mutants lead to RNAPII accumulation around the +1, and to a lesser extent the +2, nucleosome dyad (**Figure 7A, B**), but there is no change in nucleosome positioning in these mutants compared to WT cells (Chávez et al., 2001; Gutiérrez et al., 2017). This would support a function for Spt4 in maintaining efficient transcription elongation by facilitating the movement of RNAPII through nucleosomal barriers.

In conclusion, our results corroborate structural and *in vitro* studies, that implicate Spt4 as an important factor for efficient RNAPII movement through nucleosomal barriers. Importantly, this study further reveals that the contribution of Spt4 to transcription is non-uniform across the transcription unit, but more substantial in early elongation, particularly at the +2 nucleosomal barrier. We expect that future studies will address if this function of Spt4 is conserved in mammals and if the mammalian counterpart of Spt4 has a function in the RNAPII pausing observed in early transcription.

### Limitations of study

We revealed that Spt4 promotes RNAPII movement through nucleosomal barriers in vivo, and the affinity of Spt4/5 with RNAPII is lower at the upstream face of the nucleosomal DNA and lower at the downstream, implying that Spt4/5 dynamically interact with RNAPII as it transcribes through nucleosomes. As TEF-seq was performed in bulk cultures, we cannot conclude if oscillation of Spt4/5 are due to the factors fully coming on and off RNAPII or changes in the relative affinities of the factors with RNAPII. This could be addressed in the future using single-molecule approaches including RNAPII, Spt4/5 combined with nucleosomal DNA templates.

## Acknowledgements

We thank Frank Holstege for providing the *S*.*cerevisiae* Anchor Away strains, Lidia Vasilieva for Rpb9 FLAG-tagged *S*.*pombe* strain, and Micron Oxford for imaging support. This work was supported by a Cancer Research UK (CRUK) grant (C5255/A23225), a CRUK Oxford Centre Prize DPhil Studentship to Ü.U., an EPSRC studentship to T.B. (EP/F500394/10), a Royal Society University Research Fellowship to A.A. (UF120327) and BBSRC grants to J.M. (BB/S009035/1 for A.A. and BB/P00296X/1 for H.F).

## Author Contributions

Ü.U. and J.M. designed the experiments. Ü.U. performed experiments and analysed datasets. T.B. and A.A. performed and analysed mathematical modelling. H.F. set up the initial framework and analysis of TEF-seq and performed part of the MNase-seq analysis. Ü.U. and J.M. wrote the manuscript, with input from all authors.

## MATERIAL AND METHODS

### KEY RESOURCES TABLE

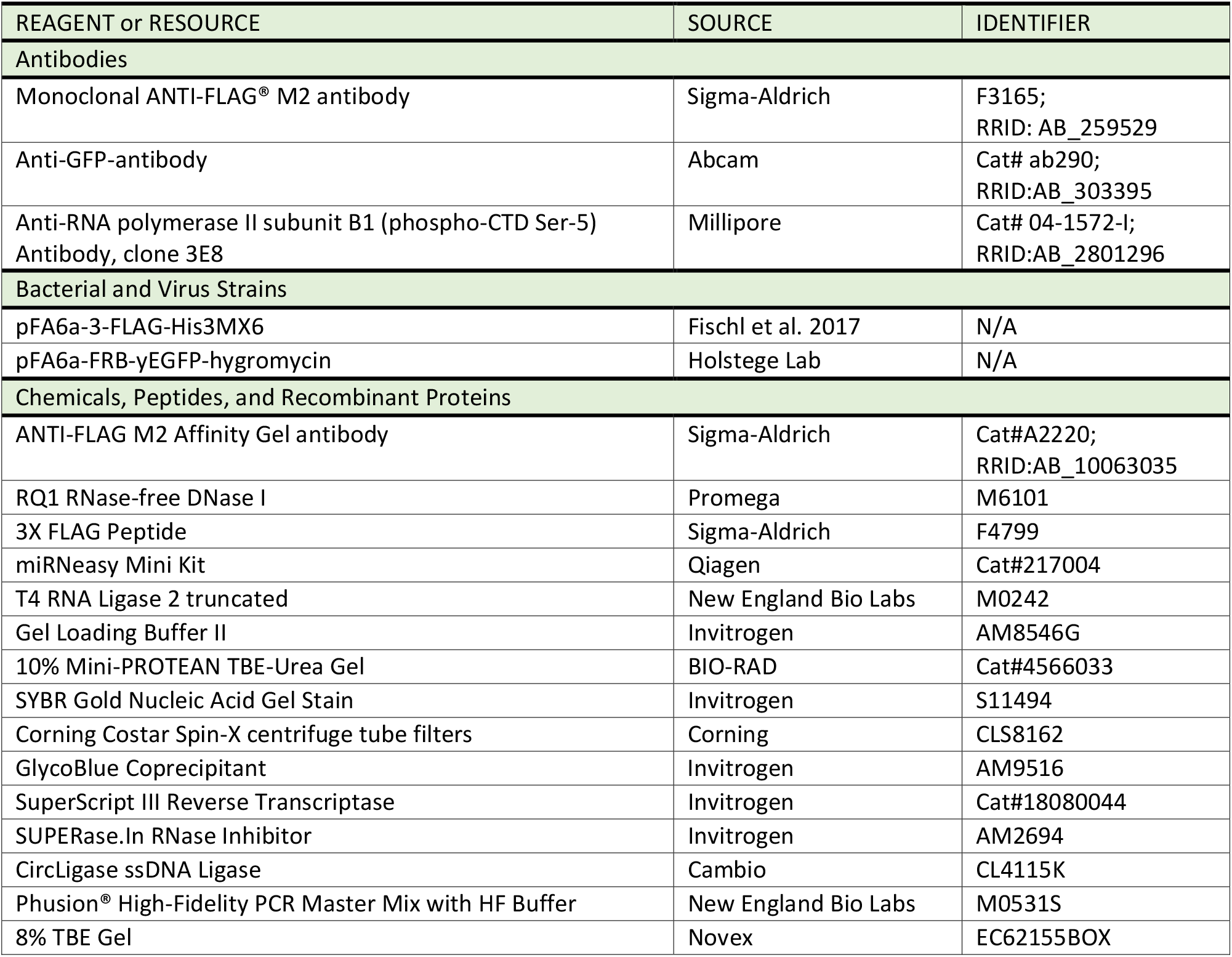
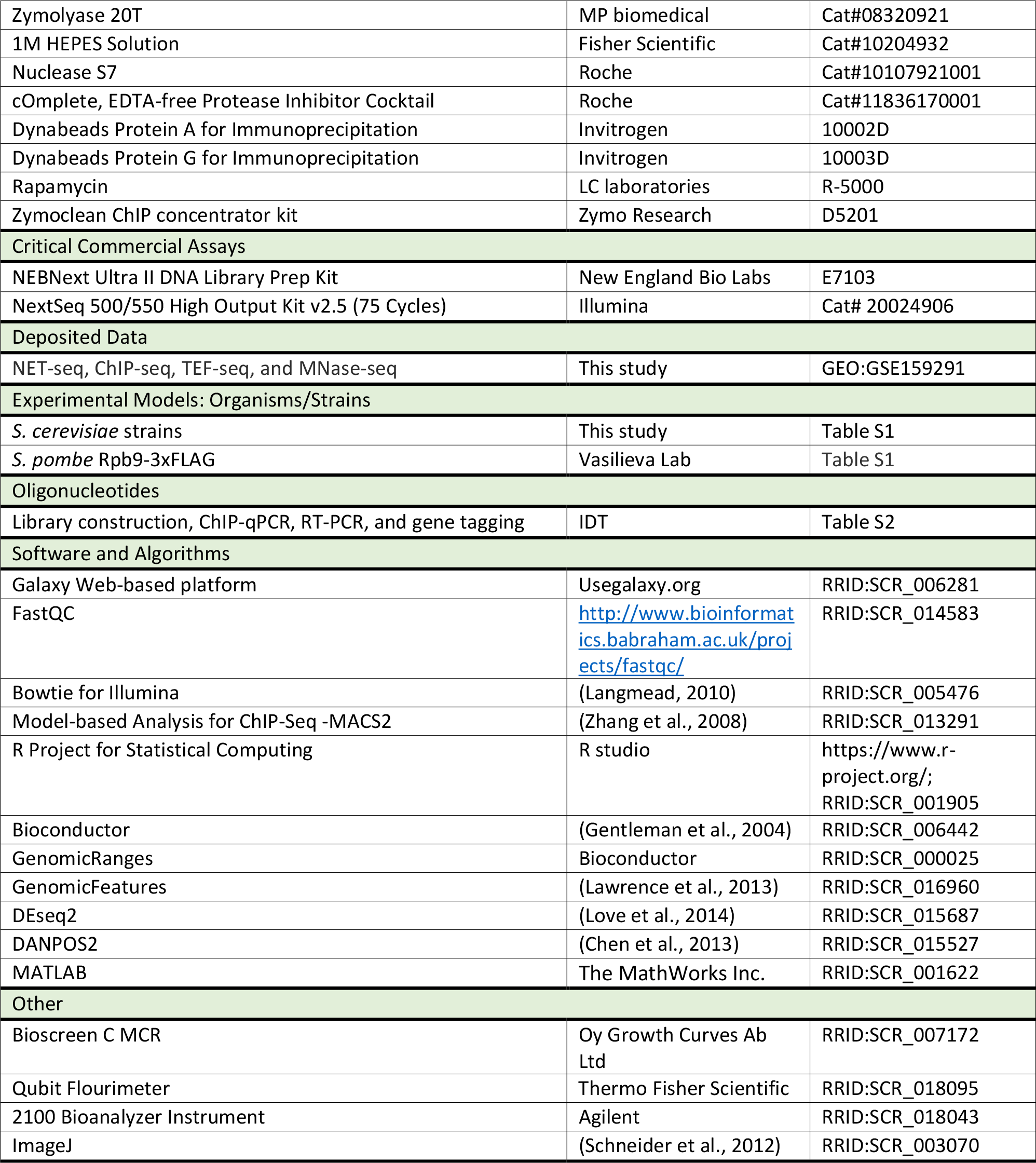

### RESOURCE AVAILIBITIY

The datasets generated during this study are available at GEO:GSE159291. The PRO-seq (GEO:GSE76142) and paf1Δ NET-seq (ArrayExpress: E-MTAB-4568) datasets were downloaded and reanalysed as part of this study. Code for mathematical model is provided at https://github.com/aangel-code/spt4_transcription_simulation.

## METHOD DETAILS

### Yeast strains and culturing

BY4741 derived *S*.*cerevisiae* cells were pre-cultured in YPD (1% yeast extract, 1% peptone, and 2% glucose) overnight at 30°C. The overnight culture was used to inoculate appropriate volume of YPD culture at OD_600_ 0.2, which was grown (30°C, 160 rpm) to OD_600_ 0.6-0.7 for all experiments unless stated otherwise. *S*.*pombe* cells were cultured in YES (0.5% yeast extract, 0.0225% of each aa: L-Adenine, L-Histidine, L-leucine, L-Lysine HCL, Uracil, and 3% glucose) in the same way as *S*.*cerevisiae* cells.

All strains used in this study, and the plasmids used to construct new strains for this study, are listed in **Table S1**. C-terminus tagging of the proteins was performed by using the homologous recombination method (Longtine et al., 1998). PCR products were amplified with a 40 bp sequence homologous to the first 40 bp upstream of the stop codon of the gene to be tagged followed by a tag sequence, selection marker and 40bp of sequence homologous to a region downstream of the gene to be tagged (see **Table S2** for primers).

### NET-seq and TEF-seq

#### Cell growth and immunoprecipitation

2 L of cells were grown in YPD to OD_600_ 0.65 (30°C, 160 rpm shaking), collected by filtering and flash frozen in liquid nitrogen. 1.28 g of frozen *S*.*cerevisiae* pellet was combined with 0.32 g of frozen *S*.*pombe* pellet. The combined pellet was ground with mixer mill (6 cycles, 3 min, 15 hz) in a metal chamber with a metal ball and the chamber was submerged into liquid nitrogen between the milling runs.

IPs were carried out in the cold room, all buffers used were ice-cold and all centrifugations were at 4°C. 1 g of grindate was resuspended in 5.66 ml of Lysis Buffer A (20 mM HEPES (pH 7.4), 110 mM KOAc, 0.5 % Triton-X-100, 0.1 % Tween 20, 10 mM MnCl2, 1x proteinase inhibitors (Roche; complete, EDTA-free), 50 U/ml SUPERase.In RNase inhibitors (Invitrogen), 132 U/ml DNase I (Promega)) by continuous pipetting up and down for several minutes. The lysate was incubated in ice for 20 min and then centrifuged (16,000 g, 10 min). The supernatant was taken and 400 µl of M2 agarose beads pre-washed twice with 10 ml Lysis Buffer A (without SUPERase.In and DNase I) was added to the supernatant. IPs were performed on a rotating wheel for 2.5 h and then washed 4 times for 2 min with 10 ml Wash Buffer A (20 mM HEPES (pH 7.4), 110 mM KOAc, 0.5 % Triton-X-100, 0.1 % Tween 20, 1 mM EDTA). Excess wash buffer was removed by centrifugation (1,000 g, 2 min). Samples were eluted twice with 300 µl 1 mg/ml of 3xFLAG peptide (Sigma) (prepared in Lysis Buffer A without SUPERase.In and DNase I) for 30 min by mild rotation. Eluates were collected by centrifugation (1,000 g, 2 min) and combined. RNAPII bound RNA was isolated with Qiagen miRNA kit according to the manufacturer’s instructions, RNA was eluted in 31 µl of elution buffer. 1 µl of the sample was used to measure RNA amount in Nanodrop. During the IP, 20 µl of samples were taken from the input, unbound (the first flow through after 2.5 h IP incubation) and eluate samples, and mixed with 20 µl of 2x SDS buffer (100 mM Tris-Cl pH 6.8, 20 % glycerol, 4 % SDS, 0.1 % bromophenol blue, 200 mM DTT) for western blot controls.

### Library preparation

#### Adapter ligation and fragmentation

A minimum of 2.5 µg of immunoprecipitated RNA was diluted in 30 µl H_2_O, split into 3 tubes and denatured (2 min, 80°C) and placed on ice (2min). RNA was ligated with 5’end adenylated and 3’end blocked adapter (**Table S2**) by adding 10 µl of ligation mix (50 ng/μl cloning linker 1, 12 % PEG 8000, 1 x T4 RNA ligase2 truncated ligation buffer, 10 U/μl T4 RNA ligase2 (truncated) (NEB)) to each tube (3 h, 37°C). Then the reaction was stopped by adding 0.7 µl of 0.5 M EDTA. Adaptor ligated RNA was fragmented by adding 20 µl of Alkaline Fragmentation Buffer (AFB; 100 mM NaCO3 (pH 9.2), 2 mM EDTA) (35-40 min, 95°C). Exact incubation time was determined for each batch of AFB. Then 0.56 ml RNA precipitation buffer (500 μl H_2_0, 60 μl 3M NaOAc (pH 5.5), 2 μl 15 mg/ml GlycoBlue (Ambion)) and 0.75 ml isopropanol was added, and samples were incubated at -20°C (>30 min). RNA was collected by centrifuge (20,000 g, 30 min,4°C) and washed with cold 80% EtOH. RNA in three tubes was resuspended in the same 10 µl of 10 mM Tris-HCl pH 7.0.

#### RNA size selection

Adapter ligated, and fragmented RNA was mixed with 10 µl gel loading bufferII (Invitrogen), denatured (2 min, 80°C) and placed on ice (3 min). Denatured RNA was run on 10% TBE-Urea gel (Biorad) (200 V, 35 min) in 1 x TBE buffer (diluted from RNase-free 10 X TBE (Ambion)). The gel was stained with SybrGold (Invitrogen) (5 min, RT) and RNA corresponding to 40-90nt was excised. For physical disruption, the gel slices were spun through 0.5 ml tubes with holes at the bottoms nested in 1.5 ml tubes. The disrupted gel slurry was incubated in 600 µl water (10 min, 70°C, 1400rpm shaking). RNA was cleared from the gel by transferring the mix into a Costar-spin column (Corning) and centrifuging (20,000 g, 3 min, RT). 50 µl 3 M Sodium Acetate (pH 5.5), 2 µl Glycoblue and 0.75 ml of isopropanol was added to RNA mix and incubated at -20°C (>30 min). RNA was collected by centrifugation (20,000 g, 30 min, 4°C), washed with 0.75 ml cold 70% EtOH and resuspended in 10 µl of 10 mM Tris pH 7.0.

#### Reverse transcription (RT)

Size selected RNA was mixed with 4.6 µl of RT mix (3.28 μl 5 x First-Strand buffer, 0.82 μl dNTPs (10 mM each), 0.5 μl 100 μM RT primer (**Table S2**)) and denatured (2 min, 80°C). Then 1.32 µl Superase.In/DTT and 0.82 µl SuperScriptIII added and incubated (30 min, 48°C). 1.8 µl 1 M NaOH added (20 min, 98°C) to degrade RNA. 1.8 µl 1M HCl added after the incubation to neutralise the cDNA.

#### cDNA size selection

cDNA was mixed with 20 µl gel loading buffer II (Invitrogen), denatured (3 min, 95°C) and placed on ice (3 min). Denatured cDNA was run on 10% TBE-Urea gel (Biorad) (200 V, 50 min) in 1xTBE buffer. The gel was stained with SybrGold (Invitrogen) (5 min, RT) and cDNA corresponding to 80-130 nt was excised. For physical disruption, the gel slices were spun through 0.5 ml tubes with holes at the bottoms nested in 1.5 ml tubes. The disrupted gel slurry was incubated in 400 µl water (10 min, 70°C, 1400 rpm shaking). cDNA was cleared from the gel by transferring the mix into a Costar-spin column (Corning) and centrifuging (20,000 g, 3 min, RT). 25 µl 3 M NaCl, 2 µl Glycoblue and 0.75 ml of isopropanol was added to the cDNA mix. Samples were incubated at -20°C (>30 min). cDNA was collected by centrifuge (20,000 g, 30 min, 4°C), washed with 0.75 ml cold 80% EtOH and resuspended in 15 µl of 10 mM Tris-HCl (pH 8.0).

#### Circularisation

4 µl circularisation mix (2 μl 10 x CircLigase buffer, 1 μl 1 mM ATP, 1 μl 50 mM MnCl_2_) and 1 µl of CircLigase (Epicentre) was added to the size selected cDNA and incubated (60 min, 60°C). Then the enzyme was heat inactivated (10 min, 80°C).

#### Amplification and barcoding

Circularised cDNA was amplified and barcoded (**Table S2**) by adding 15 µl of PCR master mix (8 µl HF master mix (NEB), 0.8 µl 10 µM reverse barcoding primer, 0.8 µl 10 µM forward barcoding primer, 5.4 µl water) per 1 µl template (1 cycle: 30 sec 98°C;; 3-to-7 cycles: 10 sec 98°C; 10 sec 60°C; 5 sec 72°C;; 1 cycle: Hold 4°C). Tubes were taken at the end of 3-4-5-7 cycles. PCR products were mixed with 3µl loading dye (NEB) and run on 8% TBE gel (Invitrogen) (90 V, 95 min) in 1xTBE buffer. The gel was stained with SybrGold (Invitrogen) (5 min, RT) and DNA corresponding to 120-170 nt was excised. For physical disruption, the gel slices were spun through 0.5 ml tubes with holes at the bottoms nested in 1.5 ml tubes. Then 0.67 ml DNA soaking buffer (0.3 M NaCl, 10 mM Tris-HCl pH 8.0, 1 mM EDTA) was added to the gel slurry and tubes were incubated overnight on a rotating wheel.

### Sequencing and data analysis

Barcoded libraries were pooled and sequenced on Illumina NextSeq 500 (50cycle, single-end) with custom reading primer (**Table S2**). Single-end FASTQ files were processed using usegalaxy.org and RStudio. Reads were groomed using FASTQ groomer for Sanger & Illumina 1.8 + (Blankenberg et al., 2010). Adapter sequence ATCTCGTATGCCGTCTTC were trimmed and reads < 15 nt were discarded using Clip function. Reads were aligned to a combined fasta file of *S*.*cerevisiae* and *S*.*pombe* genomes using Bowtie for Illumina (Langmead, 2010). SAM files were converted to BAM files using SAM-to-BAM (Li, 2011). Using RStudio/Bioconductor packages, multiply aligned reads were filtered, and reads were narrowed to the 3’ends. Selected reads were annotated to the *S*.*cerevisiae* genes derived by TIF-seq (Pelechano et al., 2013).

#### No tag normalization

NET-seq was performed on strains without a FLAG-tag to detect background signal during the IP as described in (Fischl et al., 2017). The *SCR1* gene is transcribed by RNAPIII and gives a high non-specific signal in the NET-seq and TEF-seq IPs, and this locus was used for no-tag normalisation. The reads on chrV [442007:442458] were split into 10 nt bins and FLAG-tag over no tag sample ratio is calculated for each bin. The mean *SCR1* ratio then multiplied by the no tag data and subtracted from the FLAG-tag samples.

FLAG-tag - [Mean *SCR1* ratio (FLAG-tag/no tag)] x no tag

#### Spike in normalization

NET-seq data were aligned to the combined genome of Cer3 and Pombe. After the removal of non-uniquely aligned reads and no-tag background signal, counts table was created for *S*.*pombe* genes by using RStudio/Bioconductor. Then estimateSizeFactors function in the DEseq2/RStudio package was applied to calculate the relative amounts of *S*.*pombe* reads (i.e. normalisation ratio) in each sample (Love et al., 2014). NET-seq data were calibrated by dividing *S*.*cerevisiae* reads by the normalisation ratios.

#### NET/TEF-seq metagene plots

Protein-coding genes (PCGs) >750 nt were taken and genes with negative values due to no tag normalisation were discarded. To avoid genes with wrong TSS annotation, genes having 1.5x more reads upstream of the TSS (−150 to 0 nt) than in the downstream (+1 to 150 nt) were also discarded.

PCGs were plotted relative to the TSS in a window of TSS-250 nt to TSS+750 nt or relative to the PAS in a window of PAS-250 nt to PAS+250 nt. The mean number of counts for each nt position was calculated excluding top and bottom 1% of reads to avoid random spikes introduced during sequencing. The mean number of counts then was split into 10 nt bins for the metagene plots.

### ChIP-seq

Cells were crosslinked as described in (Brown et al., 2018). Crosslinked cells were resuspended in cold FA-150 buffer (10 mM HEPES pH 7.9, 150 mM NaCl, 0.1% SDS, 0.1% sodium deoxycholate, 1% Triton X-100) and mixed with pre-crosslinked with *S*.*pombe* in 5:1 ratio (final *S*.*pombe* percentage being 16.7 %). The cell suspension was lysed with glass beads using the MagnaLyser (Roche; 6 x 45 s runs, 2500 g, 4°C). The lysate was sheared 30-40 min with a bioruptor sonicator 30 sec ON/30 sec OFF at high setting. The sheared lysate was cleared by centrifuge (10,000 g, 15 min, 4°C) and the supernatant was used for IP. 500 μl sample was incubated with ∼100 μg (25 μl) of the FLAG (M2) in 1.5 ml siliconized Eppendorf tubes for 15–20 h rotating at 4°C. When the IP was performed for ChIP-qPCR, 50 μl sample was diluted to 200 μl with FA-150 buffer and incubated with 5 μl (∼20 μg) of the GFP antibody in 1.5 ml siliconized Eppendorf tubes for 15–20 h rotating at 4°C. Bound chromatin was immunoprecipitated for 90 min at 22°C with 50 μl protein A or G-Dynabeads pre-blocked with bovine serum albumin and sonicated salmon sperm DNA. Beads and attached chromatin were pelleted, washed, and immunoprecipitated chromatin was eluted from the beads as described in (Brown, Howe et al., 2018). DNA was eluted with Zymoclean ChIP concentrator kit according to the manufacturer’s instructions. DNA concentrations were measured by qubit and libraries prepared with the NEBNext Ultra II DNA Library Prep Kit for Illumina according to the manufacturer’s instructions.

Barcoded libraries were pooled and sequenced on Illumina NextSeq 500 (75 cycle, paired). Paired FASTQ files were processed using usegalaxy.org. Illumina adapters were trimmed using Trim Galore!. Reads were aligned to SacCer3 and Pombe genomes using Bowtie2. Aligned reads were filtered to remove PCR duplicates using RmDp and filtered for quality reads MAPQ > 20 using Filter SAM or BAM. To normalise reads to Pombe spike-ins, normalisation ratio was calculated to obtain the same amount of filtered Pombe BAM reads in each sample, and SacCer3 BAM reads were calibrated using Downsample SAM/BAM accordingly. ChIP-seq peaks were obtained using MACS2 callpeak and the background signal was subtracted using MACS2 bdgcmp.

### Anchor Away – Rapamycin treatment

2.3 L of cells were grown in YPD to OD_600_ 0.3 (30°C, 160 rpm) and DMSO or 1 mg/ml rapamycin dissolved in DMSO added. For ChIP, 45 ml of cells collected at 0, 60, 140 min (or at 0, 60, and 180 min for Spt5) after rapamycin treatment. For fluorescence microscopy, 13 ml of cells collected at 0, 60, 140 min (or at 0, 60, and 180 min for Spt5).

### ChIP-qPCR

Rapamycin-treated samples were cross-linked with 1% FA (10 min, RT). Then the reaction was quenched for 5 min with the addition of 2.5 ml of 2.5 M glycine. Cells were pelleted (1,000 g, 2.5 min, 4°C) and washed twice with 10 ml cold PBS (137 mM NaCl, 2.7 mM KCl, 8 mM Na_2_HPO_4_, and 2 mM KH_2_PO_4_). Immunoprecipitation of ChIP was performed as described above. qPCR was performed using a Corbett Rotorgene and Sybr green mix (Bioline) for *RPL3* and *PGK1* loci (see **Table S2** for primers). Signal was computed using %input method.

### Immunofluorescence microscopy

Rapamycin-treated samples were kept in a falcon tube wrapped with aluminium foil to limit light exposure as much as possible. Then the harvested cells were cross-linked with 4% PFA (40 min, RT), pelleted (1,000 g, 2.5 min, 4°C) and washed twice with 5 ml of cold buffer B (1.2 M sorbitol, 100 mM KHPO_4_ pH 7.5). The pellet was resuspended in residual buffer B after the second centrifuge, and 200 µl of the suspension placed on poly-L-lysine coated coverslips and incubated (30 min, 4°C). Then coverslips were washed by dipping into MQ water twice and mounted on slides with a drop of ProLong Diamond Antifade Mountant with DAPI (Vector Shield). Slides were left at RT overnight in the dark and corners of the coverslips sealed with transparent nail polish. Slides were imaged with DeltaVision CORE wide-field fluorescence deconvolution microscope using a 100x/1.4 objective lens, T%32 filter, with exposure times of 0.05 s for DAPI and 1 s for FITC channels, respectively. For NET-seq, cells were grown to OD_600_ 0.65 (140 min for Spt4 depletion, 180 min for Spt5 depletion and 120 min for DMSO control) and 2L of cells were harvested as described above for NET-seq.

### Doubling time measurement and analysis

Overnight cultures were diluted to OD_600_ 0.10-0.15 in 250 µl YPD and grown in 100 well plates for the Bioscreen (22 h, 30°C), with readings at OD_600_ taken every 20 min with shaking (200 rpm). For the anchor away testing, YPD is supplemented with DMSO or 1 mg/ml rapamycin dissolved in DMSO. A minimum of four technical replicates were performed for each condition and strain. OD_600_measurements were analysed in R. Reads were blanked by subtracting medium-only reads. Doubling times were calculated by choosing the exponential growth phase (OD_600_ 0.2 to 0.7) and using the following equation: Doubling time = log (2)*time/ [log (max (OD_600_) – log (min (OD_600_))]

### MNase-seq

Cell nuclei was prepared as described (Almer et al., 1986).1 L of cells were grown in YPD to OD_600_ 0.6 (30°C, 160 rpm), collected by filtering and resuspended in 45 ml of cold water. Then cells were pelleted by centrifuge (1,000 g, 5min, 4°C). After discarding the water, the weight of cells (wet weight) was noted and the following volumes were used per 1 g of wet cells. 2 ml of pre-incubation solution (2.8 mM EDTA, pH 8, 0.7 M 2-mercaptoethanol) was added to wet cells and incubated (30°C, 30 min). Then samples were pelleted (1,000 g, 5 min, 4°C) and the pellet was washed with 40 ml of 1 M sorbitol. After centrifuging (1,000 g, 5 min, 4°C) and discarding sorbitol, the pellet was resuspended in 5 ml of sorbitol/B-ME (1 M sorbitol, 5 mM 2-mercaptoethanol) solution and 200 µl of 2% of zymolase solution at 30°C for 30 min with shaking. The lysate was pelleted (3,000 g, 8 min, 4°C) and the pellet was washed with 40 ml of 1 M sorbitol. Nuclei were resuspended in 7 ml of Ficoll solution (18% Ficoll, 20 mM KH_2_PO_4_, 1 mM MgCl_2_, 0.25 mM EGTA, 0.25 mM EDTA) then collected by centrifuge (20,000 g, 30 min, 4°C).

The nuclei obtained from 0.5 g equivalent of cells were resuspended in 3 ml of freshly prepared SDB (1 M sorbitol, 50 mM NaCl, 10 mM Tris-Cl pH 7.5, 5 mM MgCl_2_, 1 mM CaCl_2_, 0.075% NP-40, 1 mM 2-mercaptoethanol), split into 6 tubes. 6 reactions were set up with 20-40-80-160-320U of 10U/µl MNase (prepared in 200mM Tris pH 7.5, 50 mM NaCl, 50% glycerol; Nuclease S7 Roche) and incubated (37°C, exactly 10 min). Reactions were quenched with 50 µl of pre-warmed stopping buffer (5% SDS, 50 mM EDTA, at 65°C). 50 µl of 20 mg/ml proteinase K (Roche) added to MNase treated samples and incubated (overnight, 65°C). Samples were treated with 1µl of 10 mg/ml RNase A (1 h, 37°C) and then DNA was eluted with Zymoclean ChIP concentrator kit according to the manufacturer’s instructions. Isolated DNA was run on a 1.5% agarose-TBE gel and right digestion (80U) was chosen based on the enriched amount of mono-nucleosome bands; faint di-nucleoseome bands are still visible without over digestion. The mononucleosomal DNA band was gel extracted. DNA concentrations were measured by qubit and libraries prepared with the NEBNext Ultra II DNA Library Prep Kit for Illumina according to the manufacturer’s instructions.

### MNase-seq data analysis

Barcoded libraries were pooled and sequenced on Illumina NextSeq 500 (75 cycle, paired). Paired FASTQ files were processed using usegalaxy.org. Illumina adapters were trimmed using Trim Galore!. Reads were aligned to SacCer3 genome using Bowtie2. Aligned reads were filtered to remove PCR duplicates using RmDp and filtered for quality reads MAPQ > 20 using Filter SAM or BAM. BAM files were further analysed using a peak calling software DANPOS2 (Chen et al., 2013). Read densities and nucleosome positions obtained from DANPOS2 were used for metagene analysis.

#### MNase-seq metagene analysis

Protein coding genes (PCGs) shorter than 600 nt were discarded. Genes with 4 peaks within the first 600 nt from the TSS (+1 to 600 nt) across the three replicates were kept for avoiding genes with poorly phased nucleosomes. 2622 PCGs were left for the analysis.

### Mathematical Modelling

The process of transcription was formulated as a stochastic process with core components of initiation; elongation; polymerase occlusion; stalling; resumption of elongation from the stalled state; backtracking from a stalled state; resumption of elongation from the backtracked state; collision-induced stalling and termination; early termination with a Poisson distribution around a fixed location; two dynamic windows, in which there can be different stalling, backtracking and resumption rates; termination at the 3’ end of a gene. When polymerases collide, the situation resolves itself depending on the state of the polymerases involved. If a moving polymerase collides with another moving polymerase, the upstream polymerase becomes stalled. If a moving polymerase collides with a stalled polymerase, the upstream polymerase will become stalled and the downstream stalled polymerase will be terminated. If a moving polymerase collides with a backtracked polymerase, the upstream polymerase will become stalled.

The simulations were limited to the beginning of a synthetic gene, which covered 1000 nt. Polymerases had a fixed footprint of 40 nt. Upon reaching the end of the synthetic gene and elongating, polymerases were removed. 150,000 parameter sets were sampled uniformly between the maximum and minimum parameter values given in **TableS3**, via latin-hypercube sampling (McKay et al., 1979) and each of these was simulated for a population of 100,000 identical synthetic genes. Simulations were run for the equivalent of 40 minutes in increments of 0.005 minutes per time-step to allow the system to reach a steady state. The output distribution of transcriptionally-engaged polymerase for a given parameter set was then taken of the sum of the locations of polymerases at the final time-step of each of the 100,000 simulated genes.

For the purposes of fitting, simulation and experimental data were binned, with bins of size 10 nt. For the experimental NET-seq data, genes were defined via annotations derived from TIF-seq (Pelechano et al., 2013): the TSS and TTS for each gene was defined by choosing the most abundant start and end point detected in YPD. Only genes longer than 1000 nt in length were selected and, of those, only ones with total read counts in the first 1000 nt greater than the average for all initially selected genes. Experimental and simulated NET-seq data were normalised by dividing each bin by the total read counts or sampled polymerase locations in the first 1000 nt, respectively.

Simulated data was compared to NET-seq data using the Kolmogorov-Smirnov statistic (maximum of the differences between individual points of the CDF of each data set,(Massey, 1951)) as the goodness-of-fit metric. For the plots in **Figure S2**, the single best fitting simulation for each gene was used; for the parameter comparisons, the 100 best fitting simulations for each gene were used.

---

**Figure S1, related to Figure 1.**
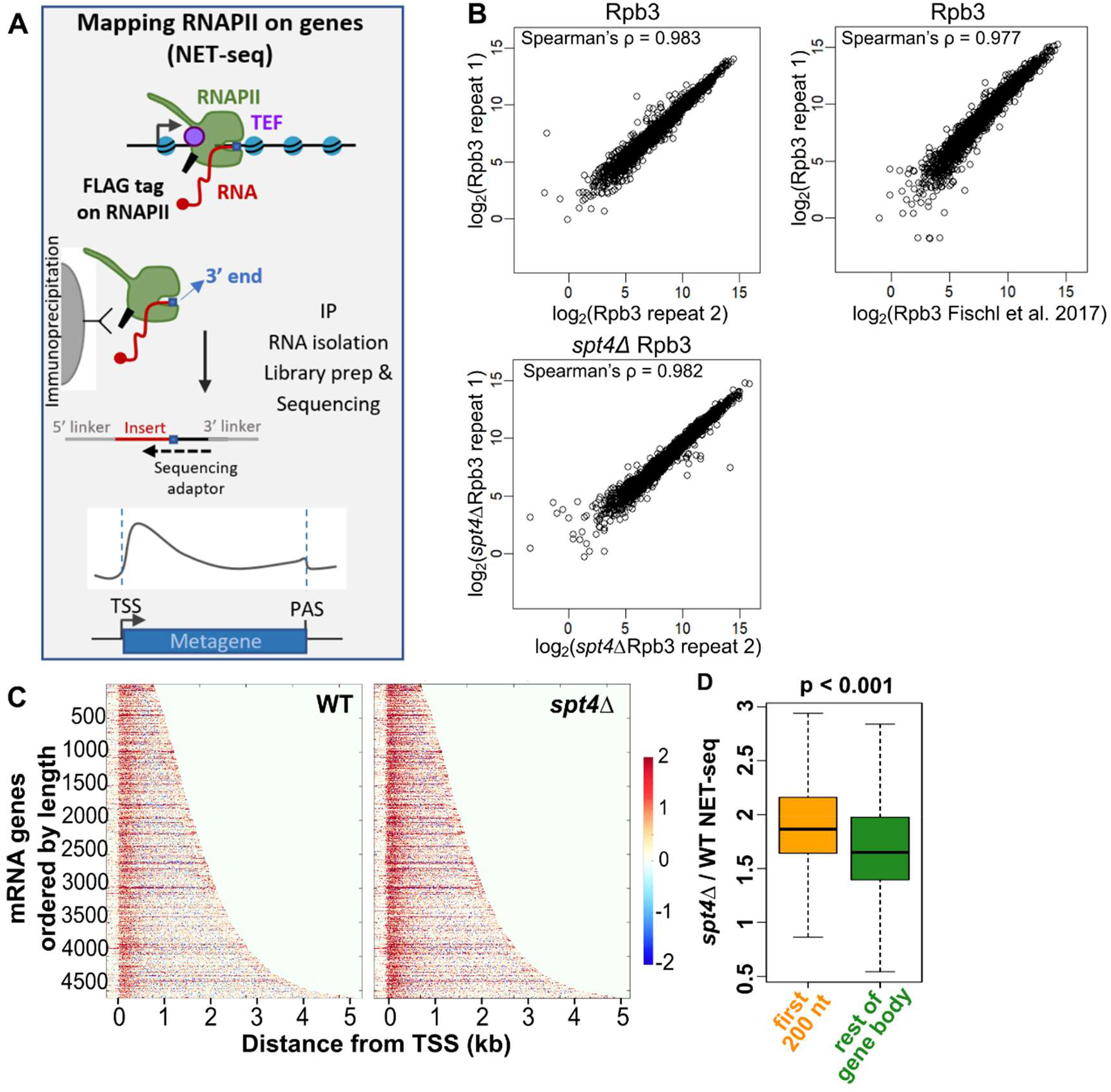
**A)** Native elongating transcript sequencing (NET-seq) pulls down elongation competent RNAPII and the 3’-end sequencing allows mapping of RNAPII at single nucleotide resolution. **B)** Correlations between NET-seq repeats from this study and with published NET-seq data from Fischl et al. (2017). Reads are counted from the TSS to the PAS for each gene. Log_2_ transformed gene counts are correlated and Spearman’s ρ calculated for each pair. **C)** Heatmaps of the WT (left) and *spt4Δ* (right) NET-seq signal. Each row indicates a PCG (n=4610), ranked by gene length. The colour code from red to blue reflects the changes in the RNAPII signal for each nucleotide position from TSS-250 nt to TSS+4750 nt (x-axis) as shown by the colour bar. **D)** Boxplots of the *spt4Δ* / WT NET-seq ratios within the first 200 nt reads from the TSS (from TSS to TSS+200 nt; orange) and the rest of the gene body (from TSS+200 to PAS-250 nt; green) for protein-coding genes after filtering low read genes out (see Methods). N=4610, p <0.001, two-tailed, paired Student’s t-test.

**Figure S2, related to Figure 2.**
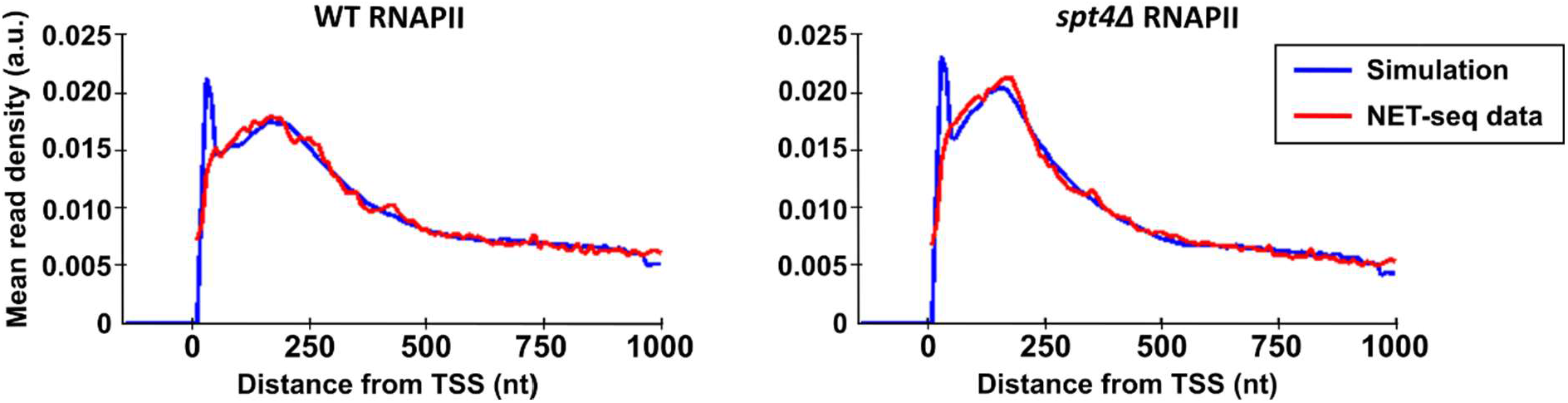
Fits of the model to the WT (left) and *spt4Δ* (right) NET-seq metagenes. Metagenes of the NET-seq data were constructed by taking the mean of the mean-normalised NET-seq reads of the first 1000 nt of reads from the transcription start site (TSS). Metagenes of the simulated data were constructed by taking the mean of the set of mean-normalised best fitting single simulation for each gene. Data are binned with a bin size of 10 nt. Notably, the early simulation peak is in a region that is not expected to be reliably detected with NET-seq protocol, therefore the difference between the simulation and NET-seq data around this region is not necessarily contradictory.

**Figure S3, related to Figure 3.**
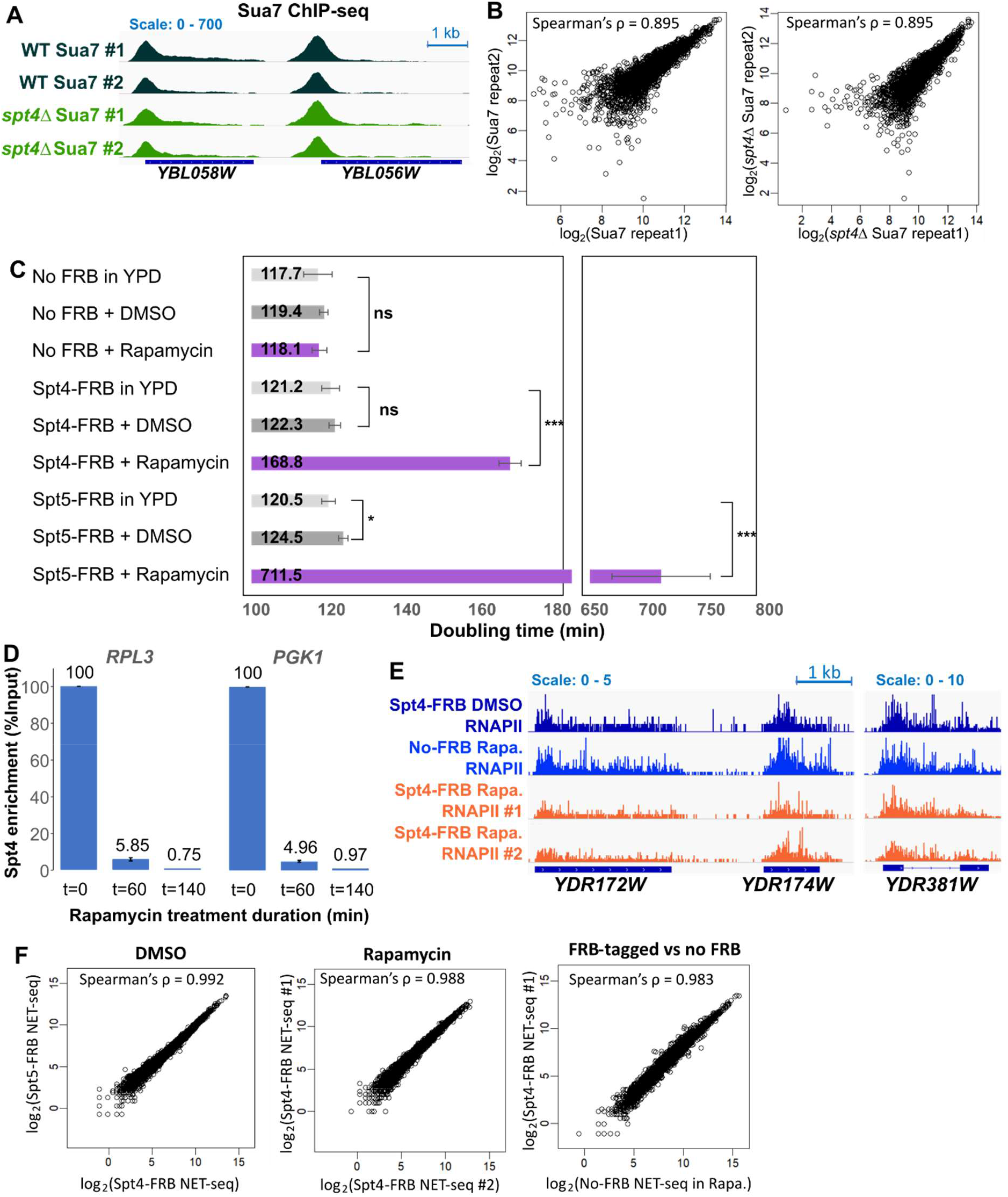
**A)** WT and *spt4Δ* Sua7 ChIP-seq signals of example genes transcribed from the positive strand: *YBL058W* and *YBL056W* in two biological replicates. The dark blue boxes indicate the transcribed region of the genes (from TSS to PAS). **B)** Correlation plot of the two repeats of each experiment. Reads are counted around the TSS (TSS-100 to TSS+100) for each gene. log_2_ transformed gene counts are correlated and Spearman’s ρ calculated for each pair. **C)** Doubling times of the anchor away strains. Cells were grown in YDP, DMSO and rapamycin (1 mg/ml in DMSO) for 22 h. OD_600_ was recorded every 20 min using the Bioscreen and doubling times were calculated for exponential growth phase (OD_600_ 0.2 to 0.8) as described in the methods. Error bars indicate standard deviation of 3 biological replicates performed at 4 technical repeats. * p-value <0.05, ***p-value <0.001 (Student’s t-test, unpaired, two-tailed). **D)** ChIP-qPCR for Spt4 upon depletion of Spt4 protein by Anchor Away across different time points. Percentage of Spt4 levels relative to time point 0 levels at the two representative genes *RPL3* and *PGK1* tested by ChIP against GFP (targeting Spt4-FRB-GFP) followed by qPCR. Error bars indicates standard deviation of the two biological replicates. **E)** DMSO or rapamycin-treated Spt4-FRB or rapamycin-treated No FRB NET-seq signals of example genes transcribed from the positive strand: *YDR172W, YDR174W* and *YDR381W*. Two biological replicates are shown for rapamycin treated cells. The dark blue boxes indicate the transcribed region of the genes (from TSS to PAS), while the blue line indicates the intronic region in *YDR381W*. **F)** Correlations between anchor away NET-seq repeats and controls. Reads are counted from the TSS to the PAS for each gene. Log_2_ transformed gene counts are correlated and Spearman’s ρ calculated for each pair.

**Figure S4, related to Figure 4.**
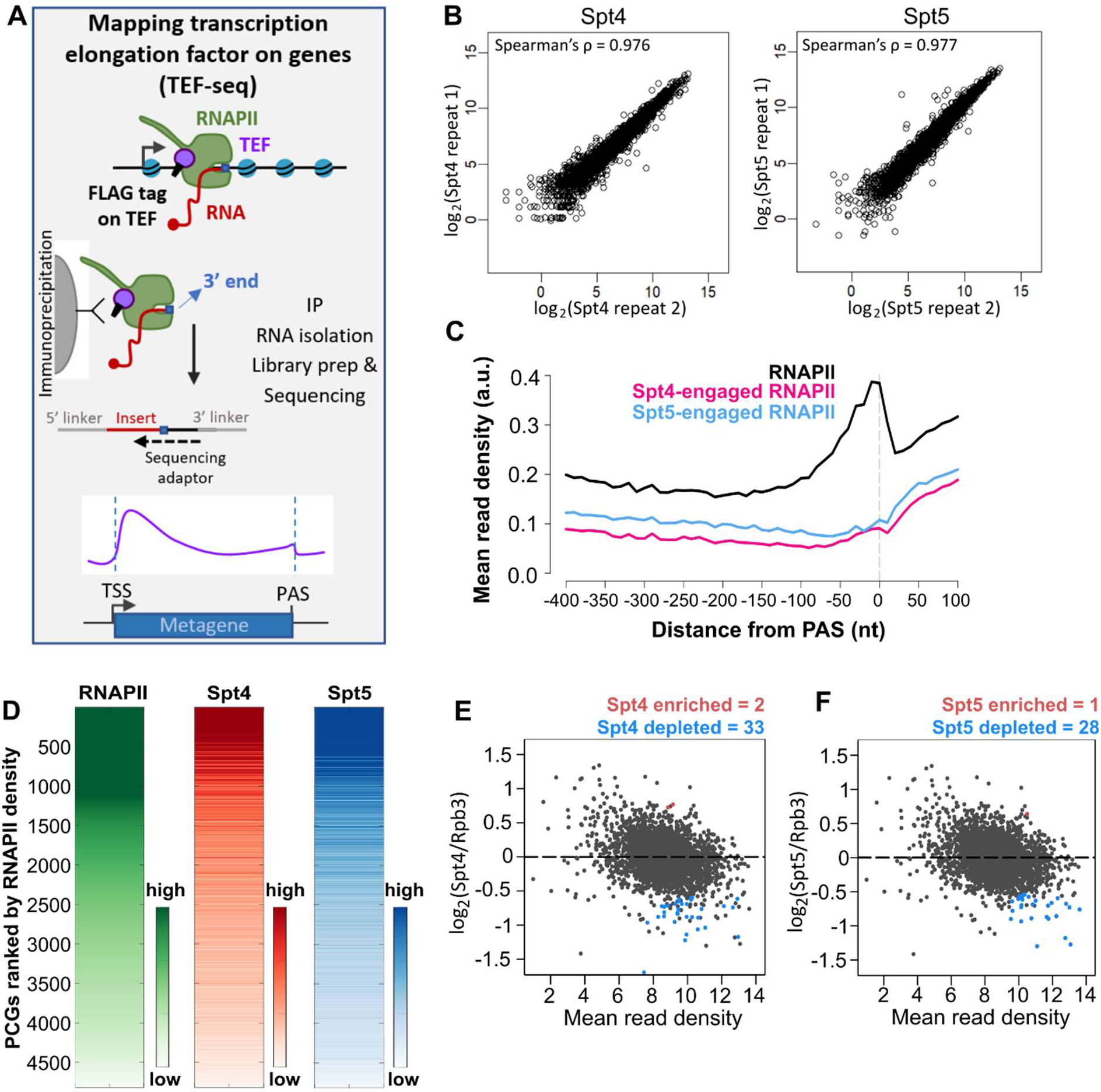
**A)** Similar to NET-seq (Figure S1A), transcription elongation factor (TEF) associated nascent elongating transcript sequencing (TEF-seq) pulls down elongation competent RNAPII from FLAG-tagged TEF. The 3’-end sequencing allows mapping of. maps TEF-associated RNAPII at single nucleotide resolution. **B)** Correlations between TEF-seq repeats. Reads are counted from the TSS to the PAS for each gene. Log_2_ transformed gene counts are correlated and Spearman’s ρ calculated for each pair. **C)** Metagene plots of NET-seq (RNAPII; black), and TEF-seq (Spt4; pink, Spt5; light blue) reads around PAS. (Close up version of Figure 4B). **D)** Heatmaps of RNAPII NET-seq and Spt4/5 TEF-seq reads over the gene bodies (taken as TSS to PAS-250 nt) on log_2_ scale. Protein-coding genes are ranked by RNAPII levels. **E) and F)** Differential enrichment of Spt4 (D) and Spt5 (E) on RNAPII. DEseq2 applied to the read counts from the gene body (TSS to TSS-250 nt) for two replicates of each data. Significantly enriched and depleted genes indicated in red and blue, respectively (p-adjusted <0.05).

**Figure S5, related to Figure 5.**
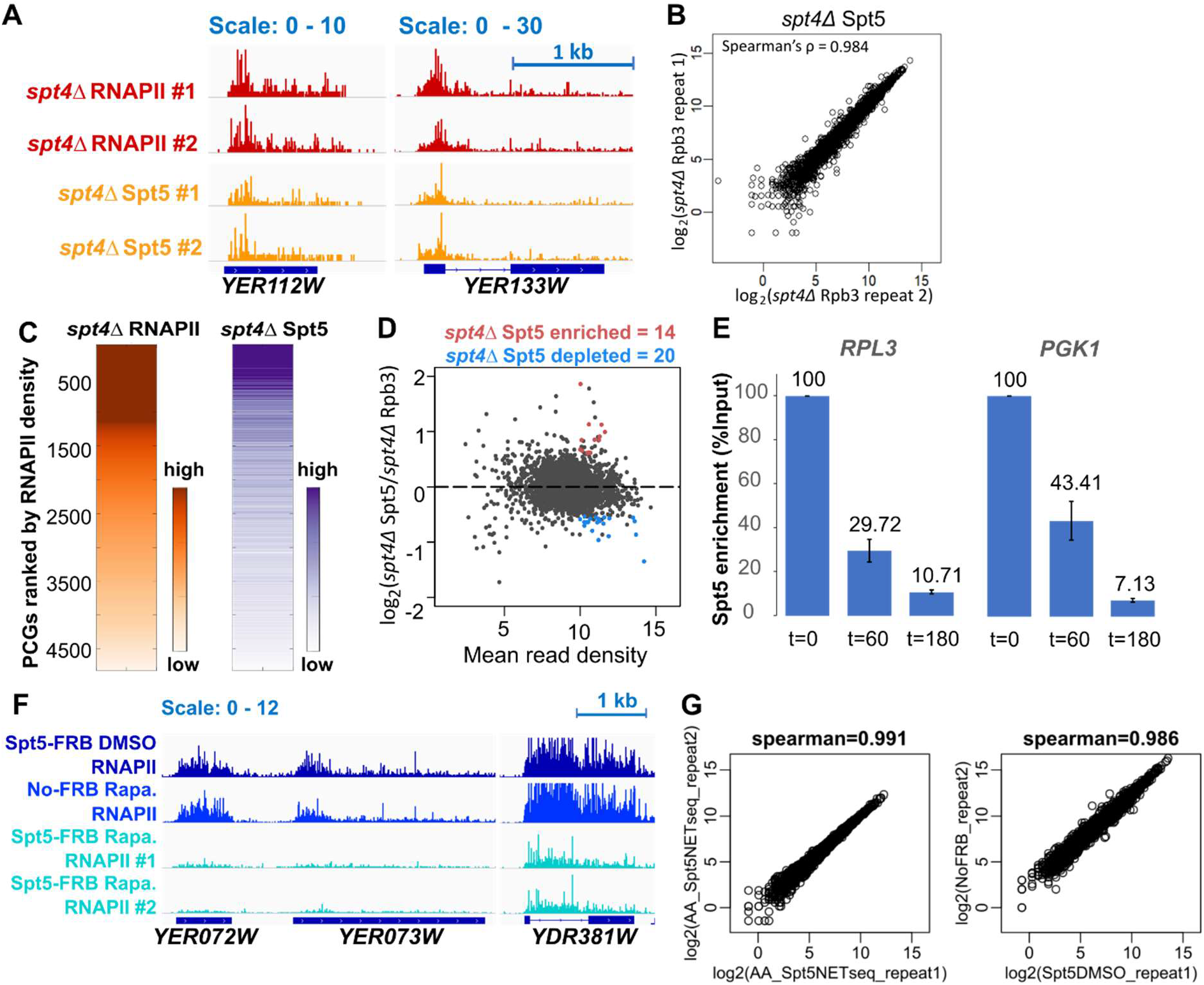
,**A)** *spt4Δ* NET-seq (RNAPII) and *spt4Δ* Spt5 TEF-seq reads of example genes transcribed from the positive strand: *YER112W* and *YER113W* in two biological replicates. The dark blue boxes indicate the transcribed region of the genes (from TSS to PAS), while the blue line indicates the intronic region in *YER113W*. **B)** Correlations between *spt4Δ* Spt5 TEF-seq repeats. Reads are counted from the TSS to the PAS for each gene. Log_2_ transformed gene counts are correlated and Spearman’s ρ calculated for each pair. **C)** Heatmaps of *spt4Δ* RNAPII NET-seq and *spt4Δ* Spt5 TEF-seq reads over the gene bodies (taken as TSS to PAS-250 nt) on log_2_ scale. Protein-coding genes are ranked by RNAPII levels. **D)** Differential enrichment of *spt4Δ* Spt5 on *spt4Δ* RNAPII. DEseq2 applied to the read counts from the gene body (TSS to TSS-250 nt) for two replicates of each data. Significantly enriched and depleted genes indicated in red and blue, respectively (p-adjusted <0.05). **E)** ChIP-qPCR for Spt5 upon depletion of Spt5 protein by Anchor Away across different time points. Percentage of Spt5 levels relative to time point 0 levels at the two representative genes *RPL3* and *PGK1* tested by ChIP against GFP (targeting Spt5-FRB-GFP) followed by qPCR. Error bars indicates standard deviation of the two biological replicates. **F)** DMSO or rapamycin-treated Spt5-FRB or rapamycin treated No FRB NET-seq signals of example genes transcribed from the positive strand: *YER072W, YER073W* and *YDR381W*. Two biological replicates are shown for rapamycin treated cells. The dark blue boxes indicate the transcribed region of the genes (from TSS to PAS), while the blue line indicates the intronic region in *YDR381W*. **G)** Correlations between anchor away NET-seq repeats and controls. Reads are counted from the TSS to the PAS for each gene. Log_2_ transformed gene counts are correlated and Spearman’s ρ calculated for each pair.

**Figure S6, related to Figure 4 and 6.**
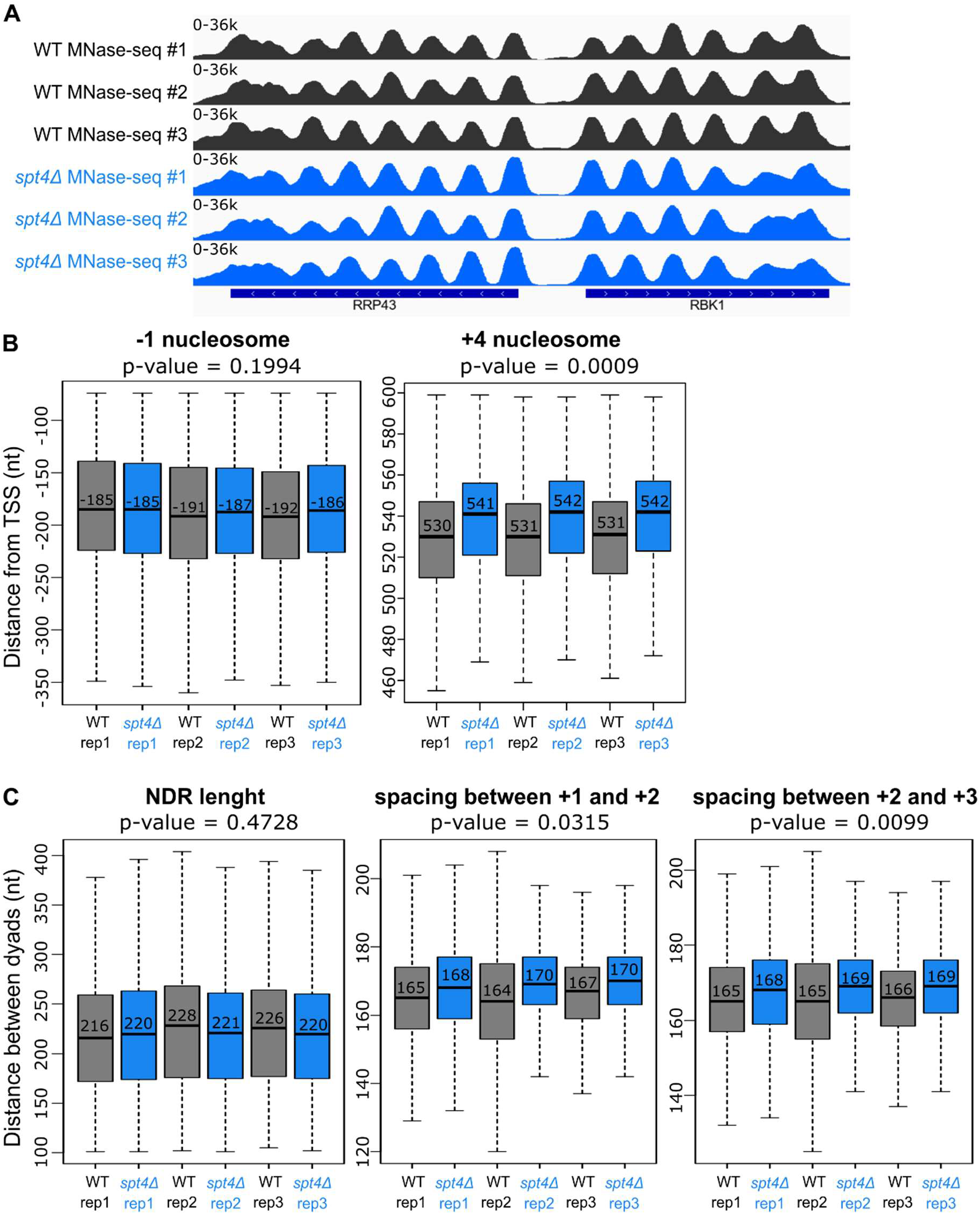
**A)** MNase-seq reads of example genes transcribed from the negative (*RRP43*) and positive strand (*RBK1*) in WT and *spt4Δ* in 3 biological replicates. The dark blue boxes indicate the transcribed region of the genes (from TSS to PAS), while the white arrows indicate transcription direction. **B)** Box-plots of the distance of the -1 and +4 nucleosomes from the TSS in three biological replicates of WT (black) and *spt4Δ* cells (blue). Numbers in the boxes indicate the median position of the given nucleosome. p-values were calculated by comparing the median position of the -1 or +4 nucleosomes in WT and *spt4Δ* conditions obtained from each replicate (Student’s t-test, paired, two sided). **C)** Box-plots of the distance between the -1 and +1 (NEDR length), +1 and +2, +2 and +3 nucleosomes in three biological replicates of WT (black) and *spt4Δ* cells (blue). Numbers in the boxes indicate the median distance between the indicated nucleosomes. p-values were calculated by comparing the median distances between the nucleosomes in WT and *spt4Δ* conditions obtained from each replicate (Student’s t-test, paired, two sided).

**Figure S7, related to Figure 7.**
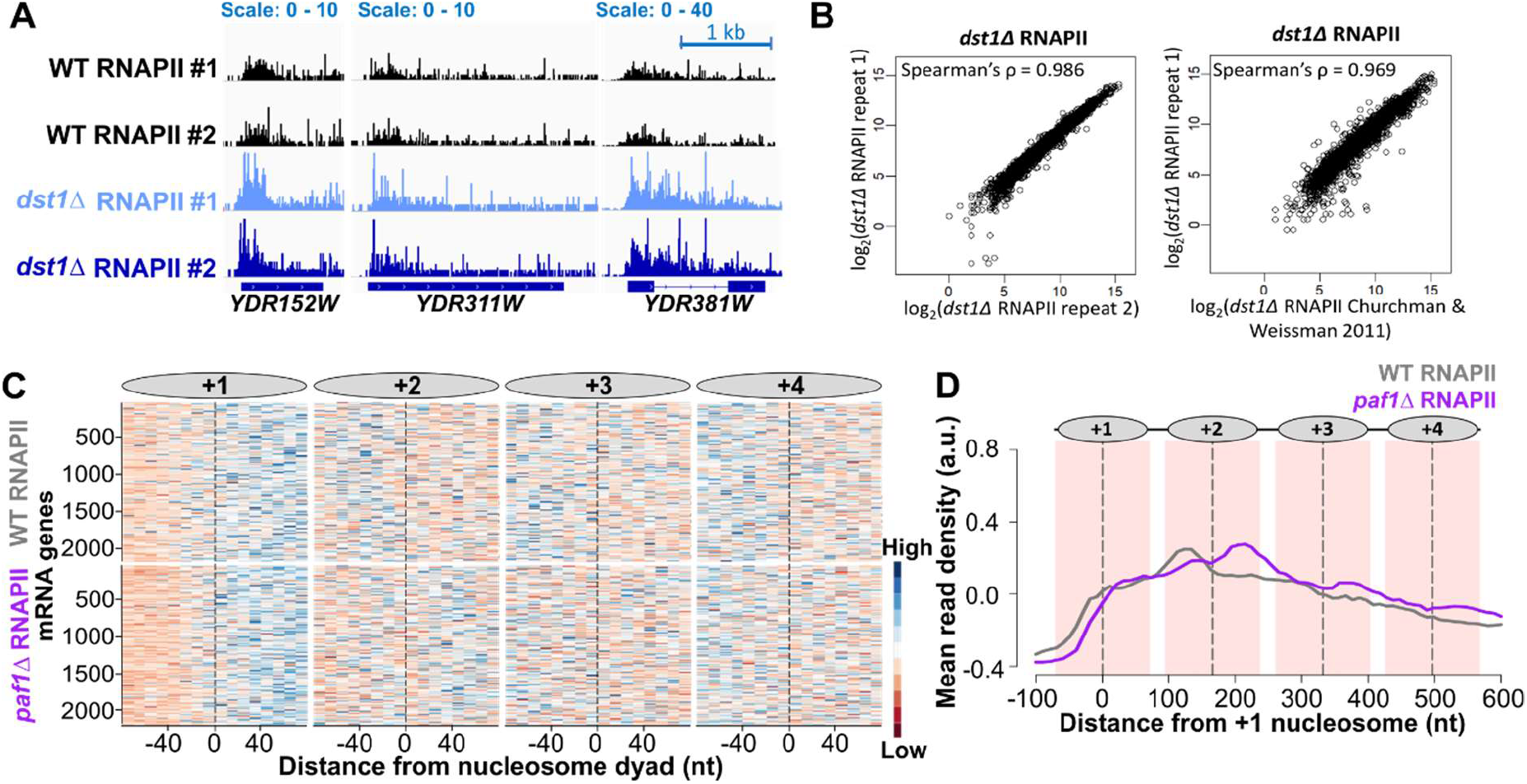
**A)** NET-seq (RNAPII) reads of example genes transcribed from the positive strand: *YDR152W, YDR331W*, and *YDR381W* in two biological replicates in WT and *dst1Δ* cells. The dark blue boxes indicate the transcribed region of the genes (from TSS to PAS), while the blue line indicates the intronic region in *YDR381W*. **B)** Correlations between NET-seq repeats in *dst1Δ* cells from this study and with published NET-seq data from Churchman & Weismann (2011). Reads are counted from the TSS to the PAS for each gene. Log2 transformed gene counts are correlated and Spearman’s ρ calculated for each pair. **C)** Heatmaps of WT (top) and *paf1Δ* (bottom) NET-seq profiles around the +1, +2, +3 and, +4 nucleosomes. Each row indicates a PCG (n=2212). RNAPII signal is shown in 10 nt bins around the indicated nucleosome dyads (−/+ 80 nt from the dyad; x-axis). The NET-seq reads were normalised to the mean and standard deviation of each gene, so that the shape of the distribution of RNAPII could be seen more clearly regardless of the expression level differences between the genes. NET-seq datasets were taken from ArrayExpress: E-MTAB-4568 (Fischl et al., 2017). **D)** Metagene plots of WT (grey) and *paf1Δ* (purple) NET-seq profiles relative to the +1 nucleosome dyad of the same data as in C. The mean and standard deviation normalised reads were used for plotting metagene profiles, as the global comparison of the NET-seq levels were not available for these datasets. Dashed lines (black) through the peaks indicate the centres of the nucleosomes and the nucleosomal DNA (+/- 70 nt around the centre) is highlighted in light pink. Position of nucleosomes graphically shown above the metagene plot.

## SUPLEMENTARY TABLES

**Table S1.** Yeast strains used in this study

| Strain | Source | Genotype |
| --- | --- | --- |
| BY4741 | Euroscarf | *MATa; his3Δ1; leu2Δ0; met15Δ0; ura3Δ0* |
| BY4741 *spt4::KanMX6* (*SPT4* KO) | Euroscarf | *spt4::KanMX6* |
| BY4741 Rpb3-FLAG (WT) | Fischl et al. 2017 | *RPB3-3xFLAG-His3MX6* |
| BY4741 *SPT4* KO Rpb3-FLAG (*spt4Δ*) | This study | *RPB3-3xFLAG-His3MX6; spt4::KanMX6* |
| BY4741 Spt4-FLAG | This study | *SPT4-3xFLAG-His3MX6* |
| BY4741 Spt5-FLAG | This study | *SPT5-3xFLAG-His3MX6* |
| BY4741 Sua7-FLAG | This study | *SUA7-3xFLAG-His3MX6* |
| BY4741 *SPT4* KO Sua7-FLAG | This study | *SUA7-3xFLAG-His3MX6; spt4::KanMX6* |
| S.pombe Rpb9-FLAG | L. Vasileva | *u+; leu1-32; ura4Δ18; ade16-M216; his3Δ1; RPB9-3xFLAG-KanMX4* |
| AA Spt4-FRB-GFP Rpb3-FLAG | This study | *tor1-1; Δfpr1; RPL13-2xFKBP12-NATMX6; met15; LYS2; his3-1; leu2; ura3; MATa; SPT4-FRB-eGFP-HygMX* |
| AA Spt5-FRB-GFP Rpb3-FLAG | This study | *tor1-1; Δfpr1; RPL13-2xFKBP12-NATMX6; met15; LYS2; his3-1; leu2; ura3; MATa; SPT5-FRB-eGFP-HygMX* |
| AA Rpb3-FLAG (No FRB) | This study | *tor1-1; Δfpr1; RPL13-2xFKBP12-NATMX6; met15; LYS2; his3-1; leu2; ura3; MATa* |
| BY4741 *dst1::kanMX6* | Euroscarf | *dst1::kanMX6* |
| BY4741 *dst1::kanMX6* Rpb3-FLAG | This study | *RPB3-3xFLAG-His3MX6; dst1::kanMX6* |

**Table S2.**
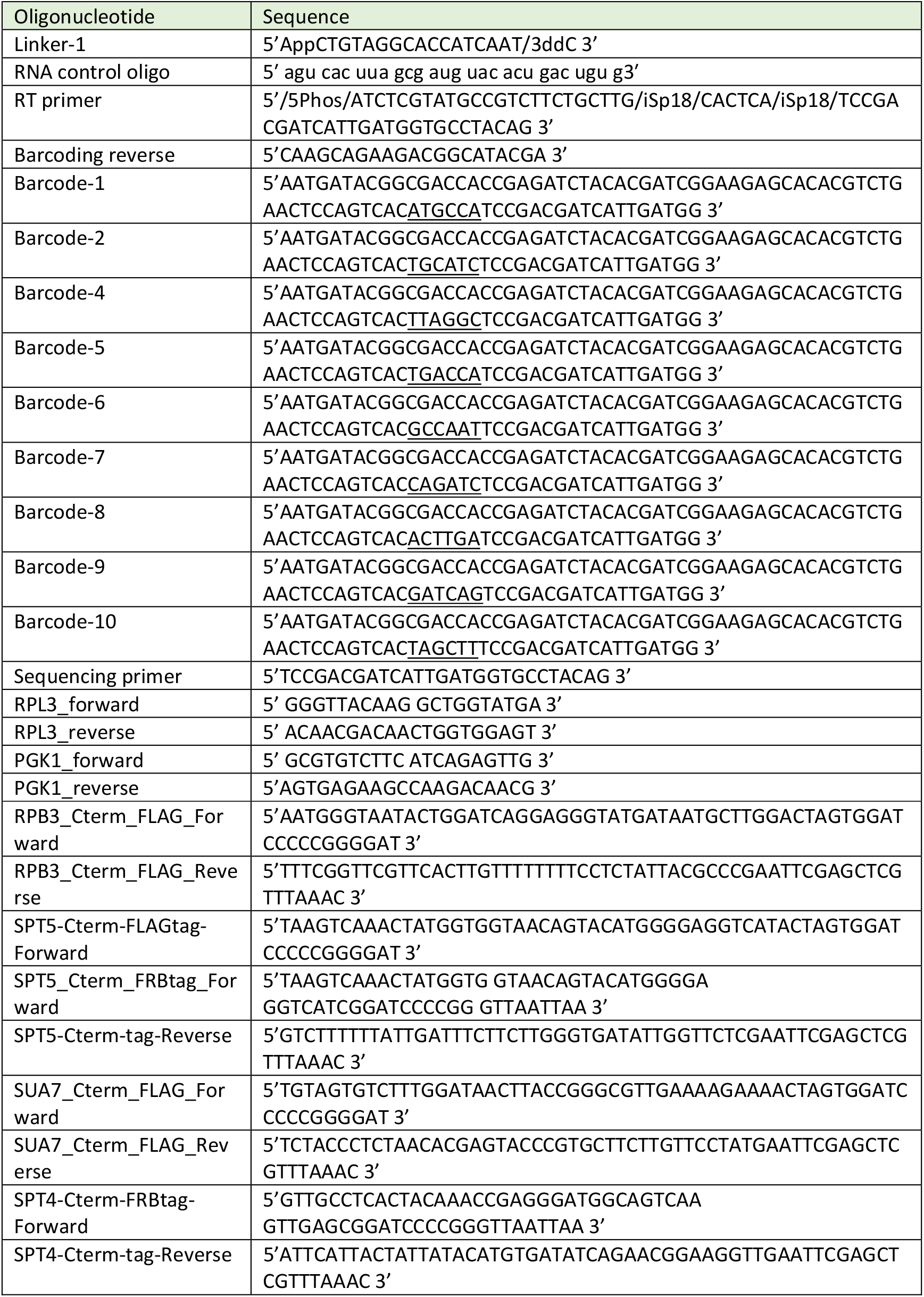
Oligonucleotides used in this study

**Table S3.** The maximum and minimum parameter values for the latin hypercube sampling

| Parameter | Min Value | Max Value |
| --- | --- | --- |
| Initiation Rate | 0.01 | 60 |
| Elongation Rate | 100 | 5000 |
| Stall 1 Rate | 0.01 | 60 |
| Stall Restart 1 Rate | 0.01 | 60 |
| Backtrack 1 Rate | 0.01 | 60 |
| Backtrack Restart 1 Rate | 0.01 | 60 |
| Stall 2 Rate | 0.01 | 60 |
| Stall Restart 2 Rate | 0.01 | 60 |
| Backtrack 2 Rate | 0.01 | 60 |
| Backtrack Restart 2 Rate | 0.01 | 60 |
| Location of Window Boundary | 0 | 1000 |
| Location of Early Termination | 0 | 1000 |

